# Factorial design as a tool to evaluate image analysis workflows systematically: its application to the filament tracing problem

**DOI:** 10.1101/806216

**Authors:** Leandro Aluisio Scholz, Ana Clara Caznok Silveira, Maura Harumi Sugai-Guérios, David Alexander Mitchell

## Abstract

The effect of image enhancement methods on the final result of image analysis workflows is often left out of discussions in scientific papers. In fact, before reaching a definitive enhancement workflow and its settings, there often is a great amount of pre-testing and parameter tweaking. In this work, we take the biofilament tracing problem and propose a systematic approach to testing and evaluating major image enhancement methods that are applied prior to execution of six filament tracing methods (APP, APP2, FarSIGHT Snake, NeuronStudio, Neutube and Rivulet2). We used a full factorial design of experiments to analyse five enhancement methods (deconvolution, background subtraction, pixel intensity normalization, Frangi vessel enhancement and smoothing) and the order in which they are applied, evaluating their effect on the signal-to-noise ratio, structural similarity index and geometric tracing scores of 3D images of a fungal mycelium and a synthetic neuronal tree. Our approach proved valuable as a tool to support the choice of enhancement and filament tracing workflow. For example, the use of deconvolution followed by median filtering gives the best geometric tracing scores if Neutube is used in the image of the fungal mycelium. Also, we show that FarSIGHT Snake and Neutube are the most robust filament tracing methods to changes in image quality. In addition, we reinforce the importance of extensive testing of new filament tracing methods against a broad range of image qualities and filament characteristics.

## 1 INTRODUCTION

Research in image processing and analysis has surged over the last two decades. Likewise, image processing and analysis are increasingly being applied in many areas, including material^1^ and life^2;3^ sciences, contributing greatly to the scientific discoveries in these areas. An increasing number of image analysis workflows is available, but this makes it ever more difficult to choose components of a workflow to solve a specific problem.

In this work, we focus on the problem of biofilament tracing, which is a common problem in bioimage analysis^4;5^, since filaments are everywhere in biology: from blood vessels and plant roots to neurons and fungal mycelia. For neuronal structures, there are many filament tracing methods from which to choose^6;7;8^. Although most of these state-of-the-art filament tracing methods were developed to trace neuronal structures^4^, this does not prevent their use with similar filament structures found in nature.

Of course, differences in image quality and filament characteristics may affect the performance of a filament tracing method, if it is used with images that are different from those for which it was initially developed. Thus, it is crucial not only to define parameters with which the methods may be evaluated, but also to evaluate the many possible combinations of methods for image enhancement and filament tracing in a systematic manner. We explore this question using a 3D image, obtained by confocal laser scanning microscopy (CLSM), of the growth of the filamentous fungus *Aspergillus niger* on agar-based media and a synthetic image that mimicks a neuronal tree.

It is essential to consider whether the quality of the image needs to be enhanced prior to filament tracing. Five of the most common types of image enhancement methods are: filament or vessel enhancement, smoothing (e.g. convolution with a median or Gaussian filter), background subtraction, image deconvolution (with a known or synthetic Point-Spread Function, PSF) and pixel intensity normalization^4^. In order to assess improvements in image quality, two parameters are commonly used: the Signal-to-noise ratio (SNR) and the Structural SIMilarity index (SSIM)^9^. The SNR carries information regarding the magnitude of the signal compared to the magnitude of the noise present in the image. Importantly, the presence and the magnitude of different categories of noise depend on the configuration of the microscopy equipment used to acquire the images. In confocal microscopy, the pinhole size, the type of detector and the scan rate all affect the SNR^10;11^.

The SSIM is one of a group of image quality parameters based on properties of the human visual system and has received special attention recently^9^. The SSIM is based on the assumption that the human visual system is highly adapted to extract structural information about objects; it considers that what one judges to be an image with poor quality results from perceived changes in structural information of the objects in the image. The SSIM is calculated as

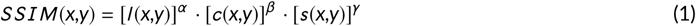

where *l* (*x,y*), *c*(*x,y*) and *s* (*x,y*) are functions that quantify luminance, contrast and structure, respectively, *x* and *y* are coordinates in the image, and *α* > 0, *β* > 0 and *γ* > 0 are weighting parameters.

Besides the image enhancement methods, we have selected six filament tracing methods with available and usable implementations: (1) All-path pruning (APP)^12^, (2) All-path pruning 2 (APP2)^13^, (3) FarSIGHT snake^14;15^, (4) NeuronStudio^16;17;18^, (5) Neutube^19;20^ and (6) Rivulet2^21^. The theoretical approaches of these methods included vary from a blend of intensity based and graph-based tracing to geometric deformable models (e.g. Multi-stencils Fast Marching). Differently than enhancement methods, improvements in tracing results may be measured by computing binary classification metrics based on geometric position comparisons if a ground truth is available: For instance, it is possible to determine the number of true positives and false negative trace points based on the distance between a point traced by the tracing method and the real (“ground truth”) position of the point^22;23^. In this context, true positives comprise segments of filament correctly traced by a method and false negatives comprise segments of filament that should have been traced but were not traced by the methods.

The aim of the present work is to demonstrate that the full factorial design is a useful tool for exploring the various possible combinations of image enhancement and filament tracing methods. We evaluated the effects of five image enhancement methods and six filament tracing implementations on two 3D images: a CLSM image of the mycelium of a fluorescent strain of a filamentous fungus and a synthetic image that resembles a CLSM image of a single neuronal tree. We used the SNR, SSIM and geometric tracing scores (e.g. recall, precision and F1-score) to evaluate the degree to which the image enhancement and tracing methods improve image quality and tracing results.

## 2 MATERIALS AND METHODS

### 2.1 Construction of the fluorescent strains

A fluorescent strain, *Aspergillus niger* pgaRed, was constructed from *Aspergillus niger* ATCC 1015 (CBS 113.46)^24^. In *Aspergillus niger* pgaRed, the expression of enhanced green fluorescent protein (eGFP) is controlled by the *gpd* promoter from *Aspergillus nidulans*, a strong constitutive promoter. The eGFP remains in the cytosol and allows visualisation of the specimen under CLSM^24^.

### 2.2 Image acquisition

*A. niger* was grown on a synthetic complete medium containing: 6.7 g.L^−1^ yeast nitrogen base for microbiology (product code 51483; Sigma-Aldrich, Germany), 20 g.L^−1^ agar, 120 mmol.L^−1^ NaH_2_PO_4_ / Na_2_HPO_4_ buffer (pH 6) supplemented with 20 g.L-1 D-Glucose^24^. Spores were spread uniformly over the solidified medium, resulting in 40 spores.mm^−2^. A small cube of the inoculated medium was excised and laid on a glass-bottom Petri dish with 4 chambers, with the inoculated surface perpendicular to the glass surface, before being transferred to the microscope. A moist cotton patch was put into one of the chambers of the glass-bottom dish to ensure that the air in the dish remained saturated with water. The inoculated medium was incubated at 30 *°*C^24^.

A confocal laser scanning microscope (Nikon A1MP+, Nikon Instruments Inc., Japan) with a temperature-controlled chamber was used to obtain 3D images at different times during growth. A 20x (0.75 NA) lens was used. The eGFP present in the fluorescent strain was excited with a 488 nm wavelength laser and detected with a filter interval (“band pass”) of 500-550 nm. Images were acquired with 16-bit grayscale bit depth and converted to 8-bit grayscale. Sample images were 973×973 px with 76 z-stacks and 1 time frame (x y z t). The *x y* resolution was 1.2361 *µ*m and the z (axial) resolution was 2 *µ*m. The sample was put under the microscope after 6 h of pre-incubation at 30 *°*C. The image stack (3D images) was acquired at 22.5 h incubation time^24^.

### 2.3 Generation of the ground truth for the image of the fungal mycelium

Several factors can make it difficult to construct a ground truth for 3D images of filaments by manual annotation: (i) large images or large numbers of images; (ii) a large number of filaments in the image or regions of high filament density; (iii) a poor image quality, for example, when photobleaching causes foreground sections of the image to be blurred or to become almost invisible. These difficulties led us to use a point annotation approach for benchmarking of the tracing methods. The approach is the same as the one used to determine the accuracy of single-particle tracking methods^23^, in which the ground truth comprises a dataset of point coordinates in an image. The procedure is represented schematically in Figure 3 (a), (c) and (d). First, the 3D image was sectioned into six subvolumes, from which three *x z* -plane images were obtained, resulting in 18 images. The *x z* -plane was used to ensure that most of the filament segments appeared as blobs in the image, given that most of the filaments grew perpendicularly to the *x z* -plane (except for the region near the surface of the agar). The *x z* -plane images were given to annotators who identified the centres of the blobs and marked them as points using the multi-point tool of ImageJ. Each of the 18 images was counted by at least three different annotators (See Supplementary Material Section 2 for more details about the procedure).

**FIGURE 1.**
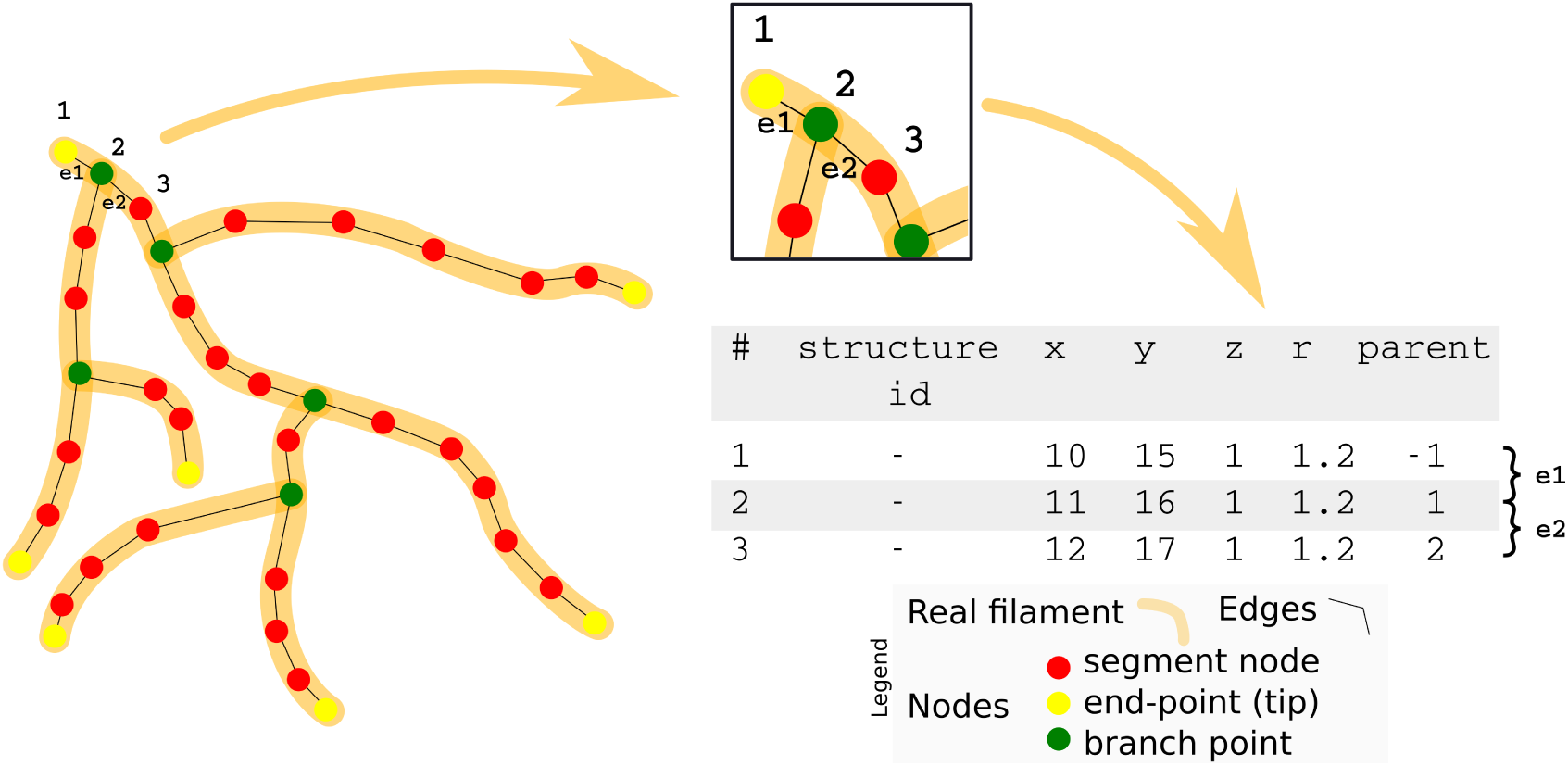
Schematic representation of the result of a filament tracing method. The nodes and edges of the tracing result should overlap with the position of the real filament. A small region with nodes (1,2 and 3) and edges (e1 and e2) is identified and shows an example output of the *swc* file format as a list of nodes that provide the node identification (id), its position in the image (*x, y* and *z* coordinates), its radius (in pixels) and the parent node, which defines the edges between the nodes.

**FIGURE 2.**
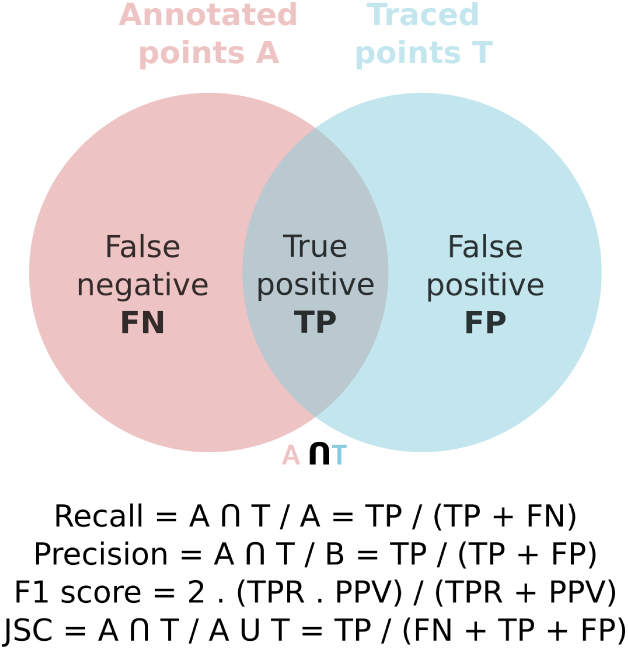
Venn diagram that shows how the four scores are calculated. *A* comprises the annotated points in the images of the *x z* plane and *T* is the point dataset of the traced images with their x,y,z coordinates. Recall is calculated by the number of matched points divided by the number of untraced plus the matched points. Precision is given by the number of matched points divided by the number of traces of non-existing filaments plus the matched points. The F1-score is calculated as the harmonic mean of the recall and precision. Finally, the JSC is calculated by the famous intersection over union calculation, which is the matched points divided by the total number of points of the tracings and annotated points.

**FIGURE 3.**
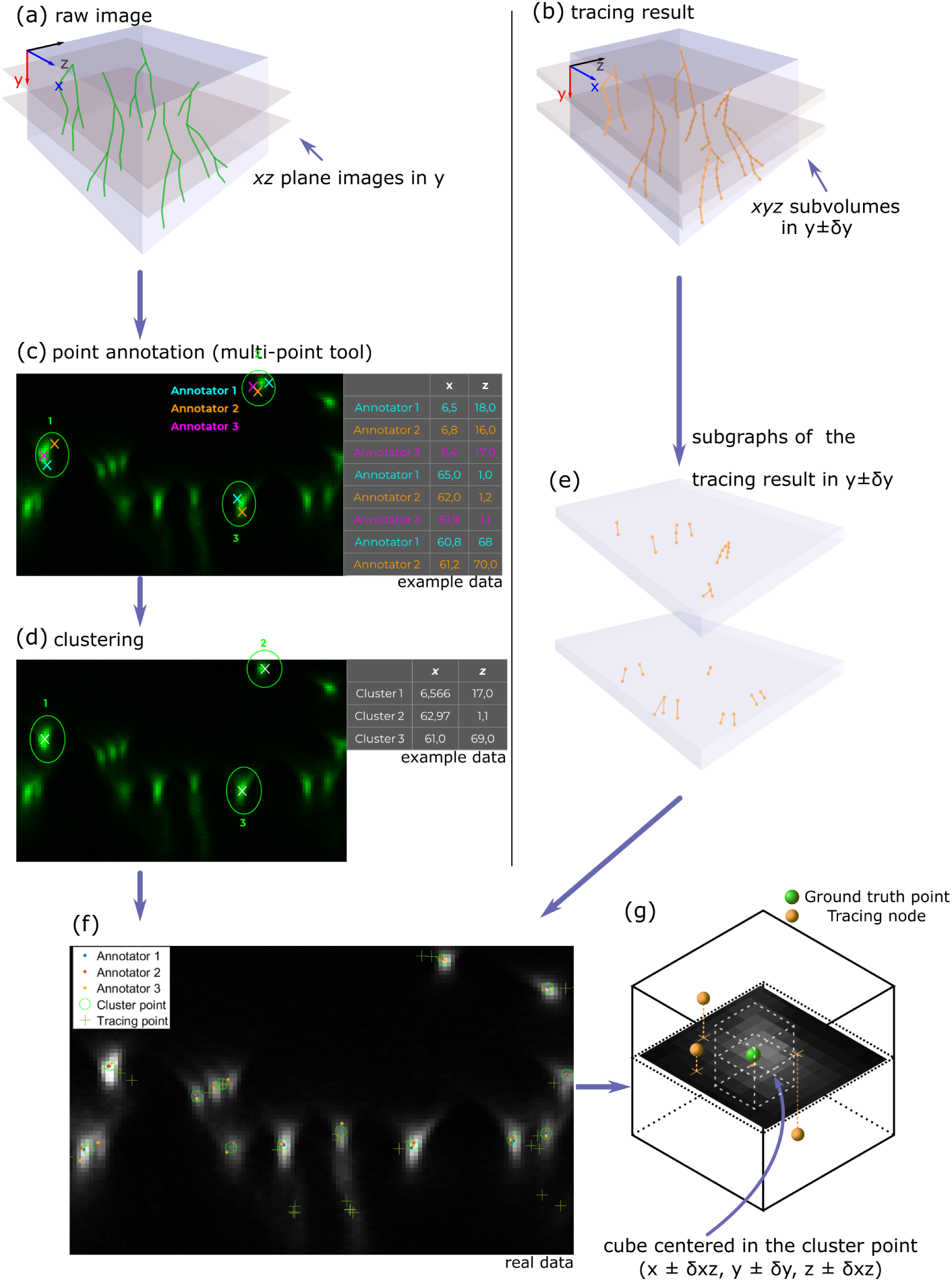
Schematic representation of the construction of the ground truth annotation and the calculation of the score. (a) representation of the raw image in three dimensions, from which images of the *x z* plane were extracted in various values of the *y* coordinate. (b) representation of the tracing result obtained from a filament tracing method. (c) sample image and example of point annotations on an image of the *x z* -plane. Each annotator determined where filament segments are located and selected a point in that region with the ImageJ multi-point tool. Then, (d) cluster points, whose coordinates are the average of a number of the closest annotated points (using euclidean distance) were determined. The maximum number of cluster points used was the maximum number of annotated points determined by any of the annotators. In order to calculate the scores, (e) subgraphs of the tracing result are extracted from the *swc* tracing file, where the minimum and maximum y coordinates of such subgraphs are determined by the parameters *δ*_*y*_ and the y coordinate of the xz plane of each annotated image as *y* ± *δ*_*y*_. Finally, the subgraphs and the cluster points are matched to calculate the scores. (f) shows an example of the data. The scores were calculated by determining a cubic region as shown in (g), where a cluster point is centered and the cube has bounds (*x* ± *δ*_*x z*_, *y* ± *δ*_*y*_, *z* ± *δ*_*x z*_). If there is at least one trace point within the region of the cube, the cluster point has a true positive tracing point, otherwise it has a false negative. The same procedure is done to the tracing points to determine the number of false positives.

The annotated points were collected and merged into a single dataset for each image. The position of each blob identified by the annotators differed slightly, hence it was necessary to use an hierarchical clustering approach in order to identify and group data points that corresponded to the same blob in the image. Thus, the maximum number of annotated points per annotator was also determined, which is a required parameter for the hierarchical clustering algorithm. This value was used as input of the clustering procedure, so that a cluster mean point could be determined. In order to obtain the cluster mean points, the Euclidean distances between each annotated point and all the remaining annotated points were calculated using the Matlab® `pdist` function. The points that belong to each cluster were obtained by using the `linkage` and the `cluster` functions. After finding the points that belong to each cluster, the mean points were calculated (mean coordinate values of the points that belong to the cluster), which were used later to calculate the scores.

### 2.4 Factorial design for the evaluation of the enhancement methods

A full factorial design was performed to evaluate the effect of five image enhancement methods on the SNR and SSIM (Eq. 1). The tests were performed using the 3D image of the fungal mycelium (See Supplementary Material for the raw image *Fungal_mycelium.tif*) and the following factors: Deconvolution (*Deconv*)^25;26^, Background subtraction (*BS*)^27^, image intensity normalization (*Norm*), Frangi vessel enhancement method (*Fra*)^28^ and smoothing with median filter (*Med*). The values of the parameters used to perform these enhancement steps are shown in Table 1 and were evaluated previously^4^. Two additional factors were added, in order to evaluate the degree to which the image quality results (SNR and SSIM) are affected by the order of application of the methods. Thus, the factors *ORD*_*median*_ and *ORD*_*Norm*_ relate to the order in which median filter and pixel intensity normalization were applied in the tests, respectively. Table 1 shows the seven factors considered in the factorial design and their levels and Figure 4 shows a reduced 2^5^ factorial design test table with coded factor levels. The experiments of the design shown in Figure 4 were done for the four possible combinations of ORD_*median*_ and *ORD*_*Norm*_ ((−1, −1); (−1, +1); (+1, −1) and (+1, +1)). An ImageJ Fiji^29^ macro script was written to execute the enhancement operations automatically.

**TABLE 1.** Factors considered in the factorial design and their levels

| Factor | Description | Low level (−1) | 1 High level (+1) |
| --- | --- | --- | --- |
| *ORD<sub>median</sub>* | Position of the median filter in the order of pre-processing | Median is applied after *Deconv* | Median is applied last |
| *ORD<sub>Norm</sub>* | Position of the normalization operation in the order of pre-processing | Normalization occurs before Frangi | Normalization occurs after Frangi |
| *Deconv* | Image deconvolution with calculated PSF<sup>1</sup> | Do not apply image deconvolution | Apply image deconvolution with calculated PSF |
| *BS* | Operation background subtraction with rolling ball algorithm<sup>2</sup> | Do not apply BS | Apply BS, ball radius 20 pixels |
| *Norm* | Operation of image intensity normalization<sup>3</sup> | Do not apply normalization | Apply normalization with 0.4% saturated pixels |
| *Fra* | Frangi vessel enhancement: multiscale Hessian based filament enhancement<sup>4</sup> | Do not apply Frangi | Apply Frangi with 5 Levels, 2 px lower and 5px upper diameter |
| *Med* | Convolve image with median filter<sup>5</sup> | Do not apply median filter | Apply median filter with kernel 3 pixels |

<sup>1</sup> The PSF was calculated using *PSF Generator* plugin<sup>25</sup>, while deconvolution was done with *DeconvolutionLab2*<sup>26</sup>.
<sup>2</sup> Background subtraction was performed using the rolling ball radius algorithm<sup>27</sup> implementation<sup>30</sup> in ImageJ.
<sup>3</sup> The built-in function *Enhance contrast* of ImageJ was used to perform this operation.
<sup>4</sup> The images enhanced by Frangi were obtained by using the *imglib2* plugin implementation in Fiji.
<sup>5</sup> The built-in ImageJ function *Filters*, 'Median' was used.

**FIGURE 4.**
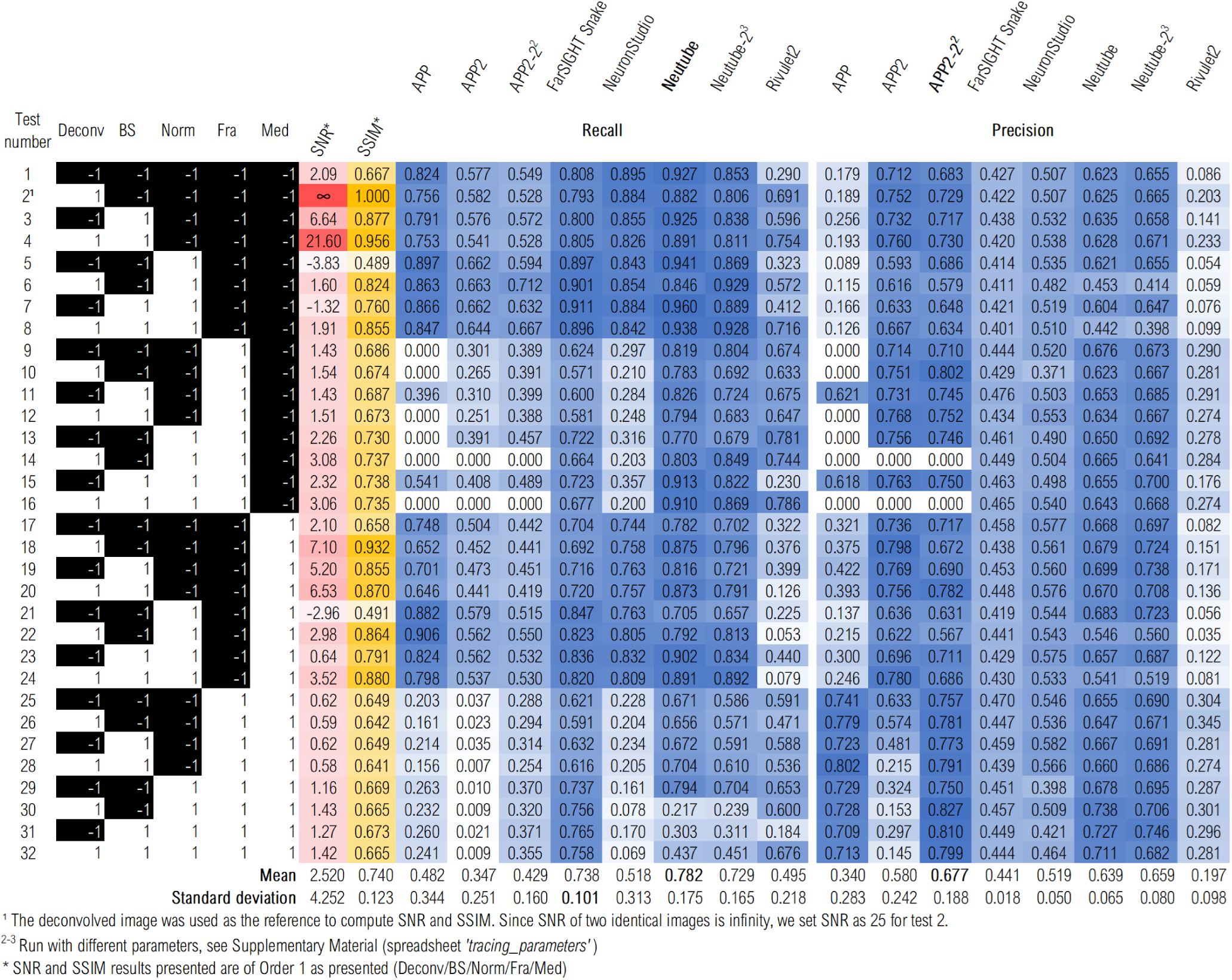
SNR, SSIM, recall and precision results of each test in the factorial design performed on the image of the fungal mycelium with the order of enhancement operations *Deconv/BS/Norm/Fra/Med*

### 2.5 Tracing of the fungal filaments

The 32 enhanced images resulting from the previous step were used as input images for six filament tracing methods: (1) All-path pruning (APP)^12^, (2) All-path pruning 2 (APP2)^13^, (3) FarSIGHT snake (FS)^14;15^, (4) NeuronStudio^16;17;18^, (5) Neutube^19;20^ and (6) Rivulet2^21^. All methods output trace results in the *swc* file format; this format comprises a graph representation of the filament tree extracted from the image. A *swc* file can only represent trees, it cannot represent filamentous structures with closed loops. Each row in the *swc* file represents a node and contains seven columns of information: the node identifier (an integer), its position in space (in Cartesian coordinates x, y, and z), a structure identifier (developed to identify neuronal structures), its radius and its parent node (Figure 1).

The filament tracing methods were then run at least once on the 32 test images (the parameter settings are available in the Supplementary Material spreadsheet *tracing_parameters.xls*). The tracing results were evaluated both quantitatively through the computation of scores (see section 2.6) and qualitatively through the visualization of the tracings. The best tracing method was selected based on two criteria: first, the score value should be one of the highest among the tested methods. Second, the connectivity of the tracing should be as accurate as possible when compared visually to the raw images of the mycelium.

### 2.6 Computation of the scores

Based on the ground truth annotations and the results of each tracing method, four different scores were computed to help evaluate the quality of the tracings: recall (or True Positive Rate), precision (or Positive Predictive Value), the F1-score and the Jaccard similarity coefficient, *JSC* (Figure 2). Single-particle tracking scores were used due to the difficulty in generating complete manual tracings of our images (the hyphae in the image were densely packed in some regions and the quality of the images made manual tracing too difficult).

The effects of the factors and their interactions on the F1-score were also evaluated. The result of the factorial design is an adjusted linear model that describes the value of the outcomes as a function of each factor and their interactions (combinations):

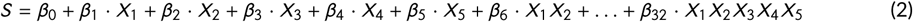

where *S* is an outcome (F1-score), *X*_1_, *X*_2_ … *X*_5_ are the coded factor levels (−1 and 1) and *β*_0_, *β*_1_ … *β*_32_ are the coefficients for the factors and their combinations.

Three parameters are required in order to calculate the scores, the spatial tolerances in the x, y and z planes around the annotated or traced points that will be considered in the point matching process, defined as *δ*_*x*_, *δ*_*y*_ and *δ*_*z*_. All tolerances are given in pixels. Figure 3 provides a detailed graphical representation and description of the calculation of the scores.

### 2.7 Comparison of the tracing results of the synthetic image with the known ground truth

The same procedures described in 2.4 and 2.5 were performed on a synthetic image generated with the TREES toolbox^31^. The resulting ground truth graph representation was converted into a 3D binary mask with Vaa3D^32^. Then, the image was convolved with a synthetic PSF (Born and Wolf) generated with PSF Generator^25^ and noise was added with the help of RandomJ^33^. Following the enhancement tests and tracing with the same methods listed in 2.5, recall, precision, F1-score and JSC were calculated for the tracing results. In this case, since a ground truth was available, the scores were computed using the complete list of nodes from the ground truth.

## 3 RESULTS

### 3.1 Enhancing and tracing the image of the fungal mycelium

We executed a series of image enhancement operations on the 3D image of the fungal mycelium and calculated the SNR (Figure 4, columns shaded red) and SSIM (Figure 4, columns shaded yellow) of the enhanced images for all 128 tests that comprise the full factorial design, which considers the factors ORDmed and ORDnorm. Figure 4 shows the results for the order DECONV/BS/NORM/FRA/MED (The results for the other orders are shown in the Supplementary Material Figures S1.1 and S1.2). The different orders did not affect the SNR and SSIM significantly: An analysis of variance (ANOVA) for the SNR gave a *p*-value of 0.939 (F value = 0.136 *α* = 0.05), while a Kruskal-Wallis test, gave a p-value of 0.2097 (*χ*^2^ = 4.52295, *α* = 0.05). Likewise, an ANOVA for the SSIM values gave a p-value of 0.971 (F value = 0.08 *α* = 0.05). Thus, we only considered the order of enhancement operations indicated in Figure 4 (see Supplementary Material Section 1 for more details). The calculation of the SNR and SSIM used the deconvolved image (test 2) as the reference image. Thus, the values of test 2 correspond to the maximum possible values for both SNR and SSIM.

The next highest values of SNR were those of tests 4, 18, 3 and 20. These tests correspond to the tests that used deconvolution and background subtraction (test 4), deconvolution and median filtering (test 18), background subtraction (test 3), and all three (test 20 included deconvolution, background subtraction and median filtering). Furthermore, two groups of tests, 1 to 8 and 17 to 24, resulted, on the whole, in relatively high SNR values; in these groups Frangi vessel enhancement was not applied to the images. However, tests 5, 7 and 21 within these groups have relatively low values of SNR. These relatively low values correspond to the use of pixel intensity normalization without prior deconvolution (5 and 7) and to the use of background subtraction, pixel intensity normalization and median filtering. The use of pixel intensity normalization increased the level of noise in the image, resulting in a SNR lower than zero dB and the prior subtraction of the background did not reduce the noise levels in tests 7 and 21. Similarly to the SNR results, after test 2, tests 4, 18 and 3 gave among the highest values of SSIM: the best SSIM values were for tests 4, 18, 24 and 3, in this order. Test 24 did not give a particularly high SNR (it was the 7th highest SNR value); it is the test in which all enhancement methods were used, except Frangi vessel enhancement. As was the case with the SNR values, the test groups 1 to 8 and 17 to 24 gave relatively high SSIM values on the whole, although tests 5 and 21 gave the lowest SSIM values.

All enhanced images were traced with six different filament tracing methods and the tracing results were compared with a ground truth in order to calculate recall and precision (Figure 4, columns shaded blue). The first set of columns shaded in blue shows the recall results and the second set shows precision results for eight runs with different sets of parameters of the six filament tracing methods. Generally, there are large differences among the values for each method. With respect to recall, the worst performing tracing methods were APP2, APP and Rivulet2. When APP2 was applied to images that had been enhanced without using Frangi vessel enhancement, the recall values were higher. On the other hand, when APP2 was applied to images that had been enhanced using Frangi vessel enhancement, the recall values were low and, in some cases, APP2 failed to trace filaments in the regions where ground truth points exist (six subvolumes of the whole image were annotated, not the entire image) thus giving recall values of zero. Also, APP did not perform well with images in which Frangi vessel enhancement had been used. The lowest values of recall were obtained for images that had been enhanced using both Frangi vessel enhancement and median filtering (tests 25 to 32). The best performing tracing methods were Neutube and FarSIGHT Snake (when recall was used as criterion), with mean recall values of 0.782, 0.729 and 0.738 (for Neutube-1, Neutube-2 and FarSIGHT Snake, respectively). For Neutube, there was a similar pattern of recall values across the tests, but the negative effect of Frangi vessel enhancement and median filtering was weaker (compare tests 1-16 with tests 17-32). With FarSIGHT Snake, the factors interacted in more complex manners: Pixel intensity normalization affected recall positively (tests 5-8, 13-16, 21-24, 29-32), whereas the use of Frangi vessel enhancement followed by median filtering had a slight positive effect compared to when Frangi vessel enhancement was applied without median filtering (compare tests 9-16 with tests 25-32).

APP2 and Neutube were the best performing methods when precision was the criterion: the highest mean precision values were obtained with APP2 and both Neutube-1 and Neutube-2 (the APP2-2 value of 0.677 being the highest). In contrast, the low precision values of the APP and Rivulet2 methods show that they oversegment the filaments, thus generating too many false positive nodes. Despite the low precision values for the majority of the tests with APP, APP had greater precision values in tests where Frangi vessel enhancement and median filter were applied together (tests 25 to 32 compared to tests 9 to 16), with the precision reaching values as high as those obtained with APP2 and Neutube. Interestingly, the effect of enhancement methods on the final precision values of all filament tracing methods was less pronounced than their effect on recall values: the standard deviation of the precision values was much lower than the standard deviation of the recall values.

Although it is valuable to analyse precision and recall results alone, it is also important that the tracing method chosen gives high values for both precision and recall. Therefore, the F1-score and JSC are better measures of overall performance of the tracing methods, since they are calculated using both precision and recall values. Figure 5 shows the F1-score and JSC for the best performing methods. The F1-score and JSC results are almost equivalent for evaluating the performance, so we focus on F1-score in the analysis that follows. APP2 was the tracing method that gave the broadest range of F1 values and also had the most uniform distribution through its range (note that the violin plot for APP2 does not show a clear peak, such as is visible in the violin plot for FarSIGHT Snake). This shows that APP2 was the method that was most sensitive to changes in the image enhancement methods used. However, some of its scores were higher than those of FarSIGHT snake, especially those scores for tests that did not include Frangi vessel enhancement and median filtering (See Supplementary Material, Table S1.2). Conversely, FarSIGHT Snake was the tracing method that was least sensitive to changes in the image enhancement methods used. This is indicated by the relatively small range of F1-values (minimum of 0.49 and maximum of 0.575) and the standard deviation of 0.022%. In the end, Neutube was the best performing tracing method when the F1-score was the criterion: in two runs with different parameters, it obtained mean F1-score values of 0.685 and 0.674 and standard deviations of 0.097 and 0.094%. The highest F1-score (0.765) was achieved with Neutube-1 in test 18, in which enhancement operations were deconvolution followed by median filtering. Neutube F1-score values were significantly better than those of the other filament tracing methods, since a Kruskal-Wallis test (*χ*^2^ = 143.13, *df* = 7, p-value= 2.2 · 10^−16^) followed by a Dunn test for pairwise comparison of Neutube against the other methods using rank sums provides the following adjusted p-values (Bonferroni method): APP = 7.403 · 10^−16^, APP2 = 5.892 · 10^−16^, APP2^2^ = 1.230 · 10^−4^, FarSIGHT Snake = 4.970 · 10^−4^ and NeuronStudio = 3.253 · 10^−5^, which confirm that the null hypothesis of the Dunn test is rejected in all cases^34^.

**FIGURE 5.**
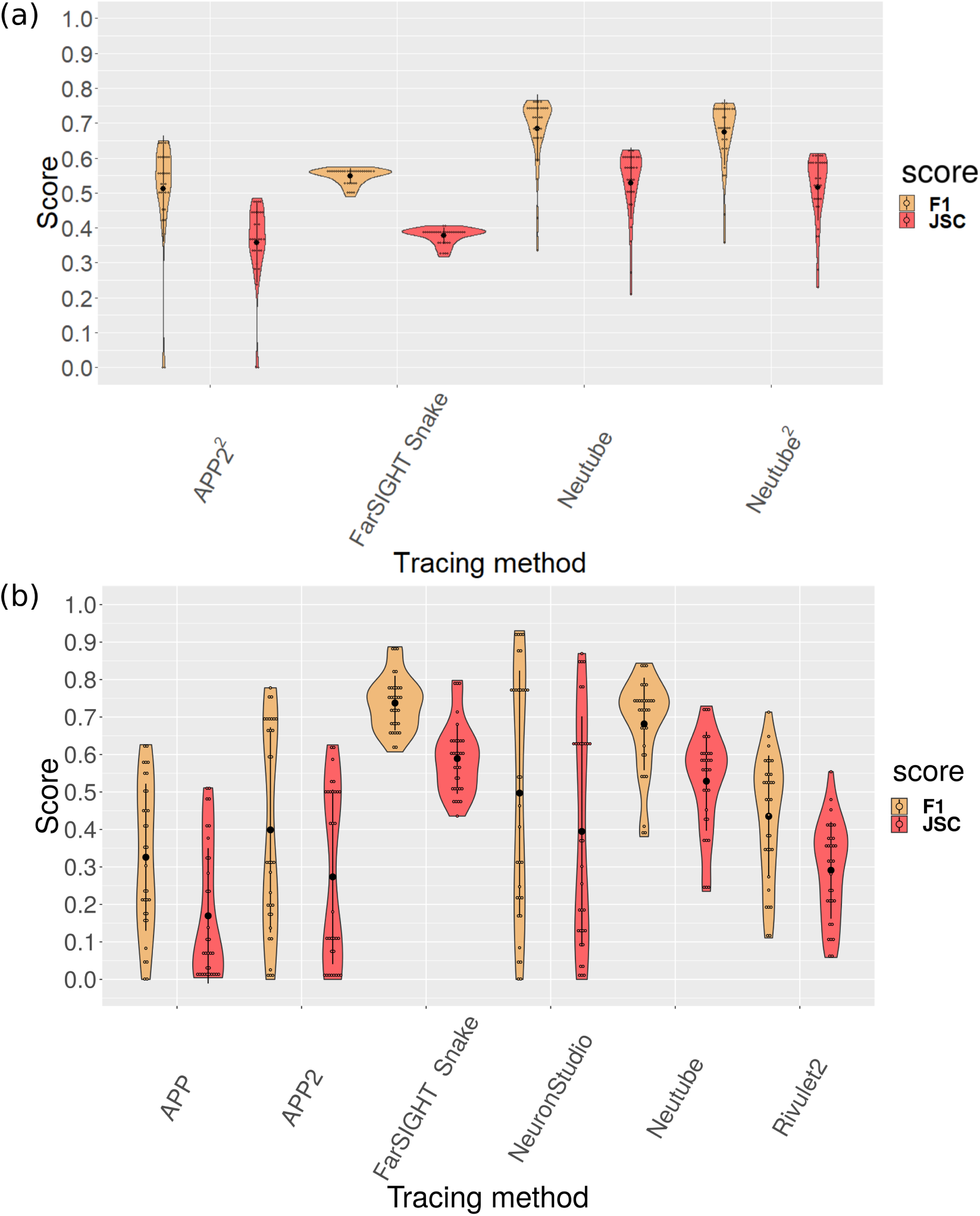
Violin plot of the F1-score and JSC values for (a) the best performing tracing methods for the fungal mycelium image: APP2^2^, FarSIGHT Snake, Neutube and Neutube^2^. (b) F1-score and JSC results of all tracing methods for the synthetic image. Empty dots are values from the 32 tests. Black dots show mean value and lines represent the standard deviation.

Figure 6 shows the test image with the results of several tracing methods. Tracing methods such as APP and Rivulet2 (Figure 6(c-d)) generally gave tracing results with too many nodes, which led to a low precision. However, a tracing result with a dense concentration of nodes (in other words, an “oversegmented” trace result) may or may not be topologically incorrect. For example, the APP results are topologically incorrect due to the spurious branches, whereas the Rivulet2 tracing results appear to be topologically correct, since the additional nodes do not form spurious branches. APP2 and Neutube gave high precision, but low recall: despite failing to segment all the filaments, they found an accurate position of the detected filaments (Figure 6(e-f)). Neutube with test 18 and NeuronStudio with test 23 were the best performing combination of enhanced images and tracing methods: They had a good balance between recall and precision (Figure 6(g-h)). This confirms that the F1-score is well suited for evaluating the results.

**FIGURE 6.**
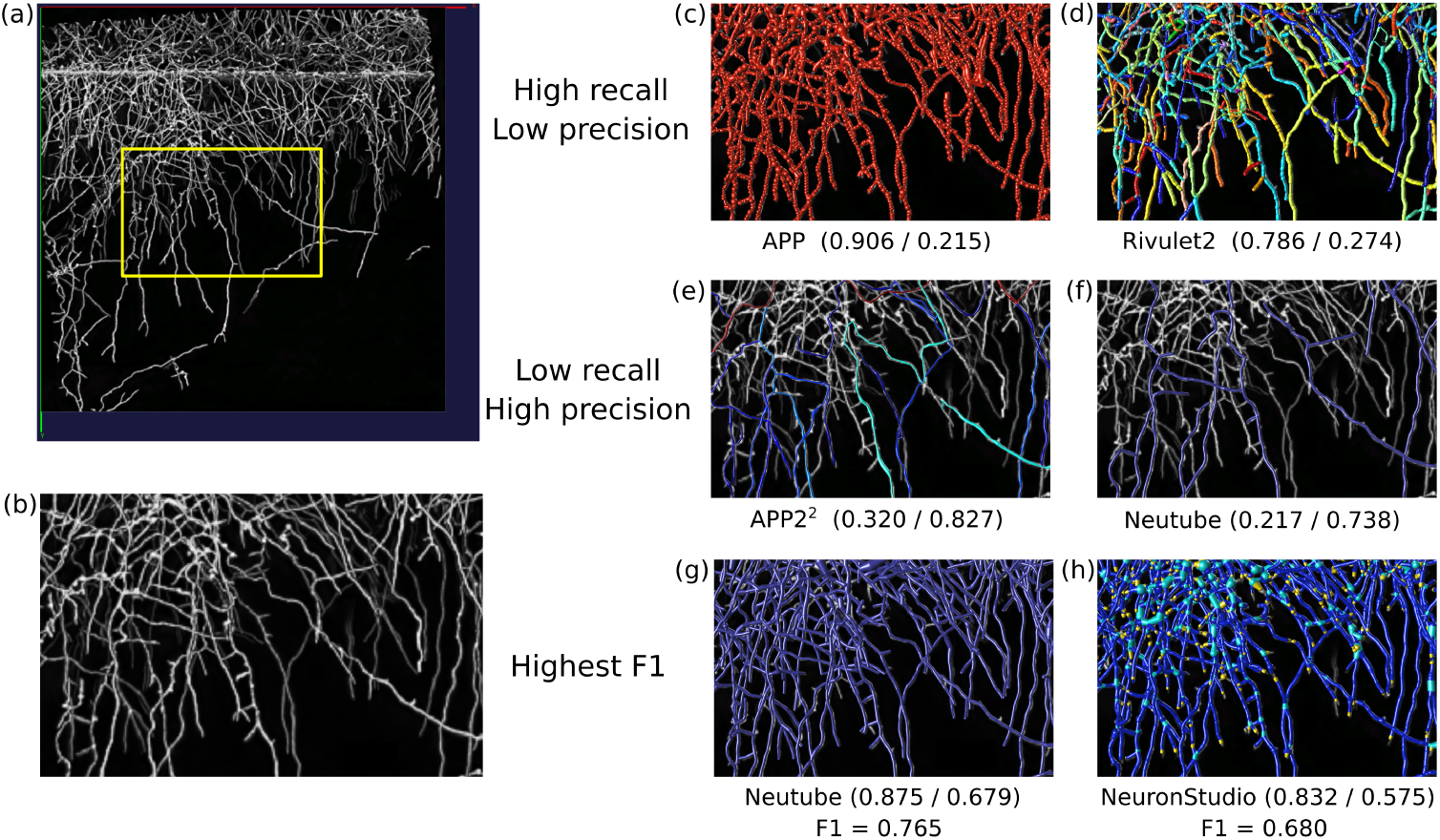
(a) 3D rendered view of the deconvolved image of the fungal mycelium (test 2), where a subregion of interest is outlined in yellow and shown in (b-h). (c-d) tracing result overlays of methods APP test 22 and Rivulet2 test 16, which gave high recall and low precision values. (e-f) tracing result overlays of methods APP^2^ test 30 and Neutube test 30, which yield low recall and high precision values and (g-h) the two best performing methods with respect to F1-score, Neutube (test 18) and NeuronStudio (test 23). Recall and precision values are provided for each tracing results within parentheses as: (recall / precision). Views were acquired with Vaa3D 3D viewer^35^.

The final outcome of the factorial design is the set of coefficients of the effects, on the F1-score, of the factors alone and in combinations. Figure 7 shows a chord graph, which represents the coefficients as chords, with the chord width corresponding to the value of the coefficient. In other words, the chord graph shows the magnitude of the effects of each factor (i.e. of each enhancement method) on the tracing score of any tracing method and also shows the overall effect of the individual factors on all tracing methods, with the magnitude of this overall effect corresponding to the arc length. For example, the most negative factor was the use of Frangi vessel enhancement alone since its arc length is the largest among the factors and most of its chords represent negative coefficient values, with APP2 and NeuronStudio having the most negative values. Deconvolution was the second most negative factor, with negative coefficients for all tracing methods except Rivulet2, although there were no significant differences in the moduli of the coefficient values of the tracing methods. On the other hand, the most positive factor was the combined use of deconvolution and median filtering but its relative importance compared to the other factors was not significant. The combined use of deconvolution, Frangi vessel enhancement and median filtering had the second most positive effect on the tracing methods.

**FIGURE 7.**
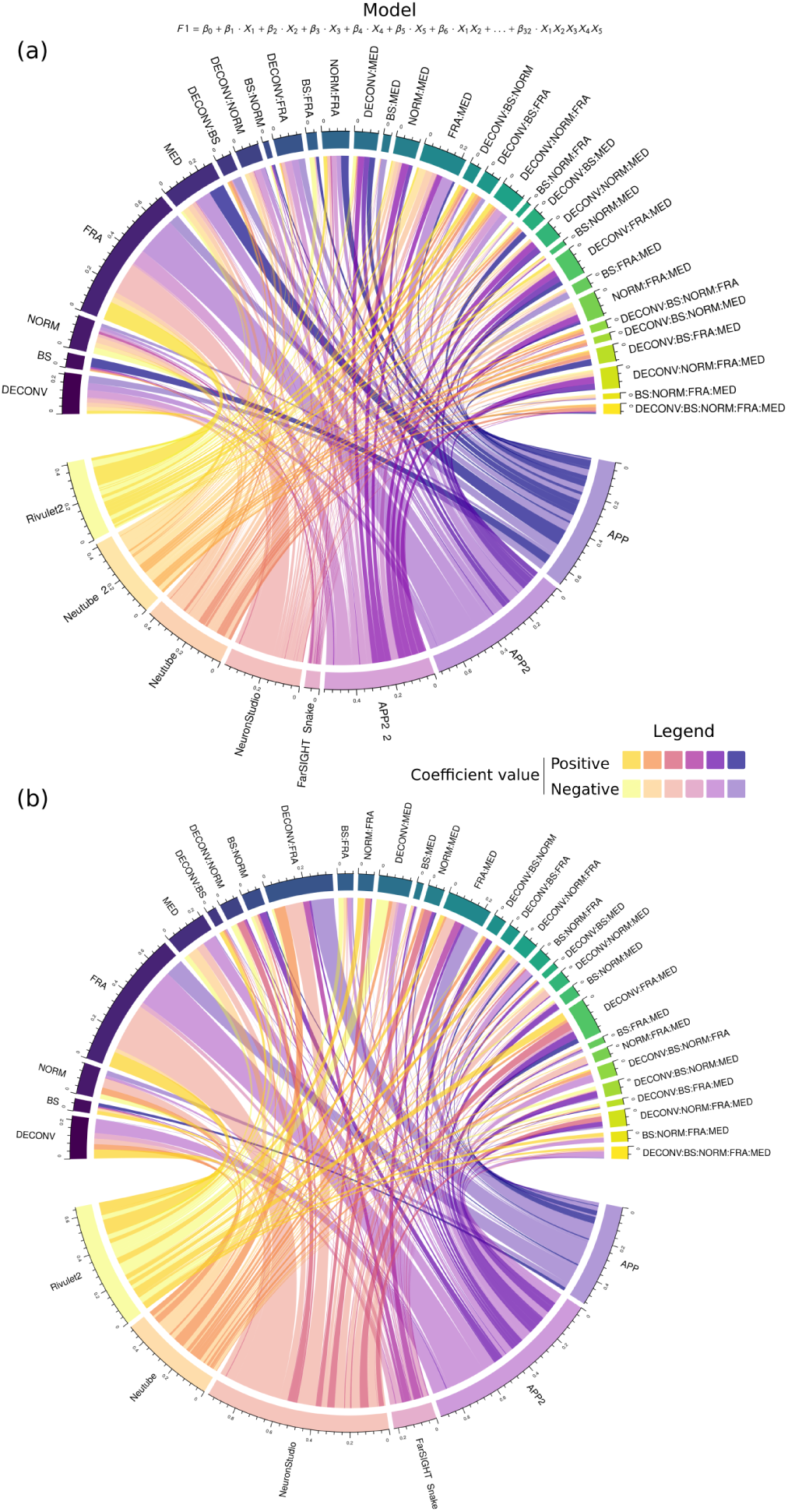
Chord graph representing the coefficients of the effects of the factors, alone and combined, on the F1 score for the tests with (a) the fungal mycelium image and (b) the synthetic image. In each graph, the tracing methods are represented by the lower arcs, the lengths of which are equivalent to the sum of the moduli of the coefficients. Thus, the length of the arc gives an idea of the spread of the F1-score values throughout the 32 tests. The model equation shown above the chord graph shows how the F1-score is modelled as a function of the factors and its combinations. The factors *X*_1_, *X*_2_, …, *X*_5_ and their combinations are represented as arcs in the upper region of the chord graph. The lengths of these arcs are equivalent to the sums of the moduli of the coefficients of each method with respect to the factor. The length of the arc of the factor is proportional to the overall effect of the factor on all tracing methods. The coefficients that multiply each factor *β*_0_, *β*_1_ … *β*_32_ are represented as links between a method and a certain factor and are depicted in two different colour tones: an opaque colour and a transparent colour (see legend). The opaque colour represents a positive value of the coefficient (i.e. a positive effect on the F1-score), whereas the transparent colour represents a negative value of the coefficient (i.e. a negative effect on the F1-score). Raw data in the form of a table is available in the Supplementary Material Table S1.1 and S1.2. Chord graphs generated with the `circlize` R library^36^.

Finally, the best method for the image of the fungal mycelium was Neutube. On the whole, the Neutube method had relatively high recall values (tests > 0.75) and intermediate precision values (tests that gave precision values between 0.5 and 0.75), an example is shown in Figure 6(g) (Neutube-1 test 18). Also, Figure 7(a) shows that there was no specific factor impacting F1-score more significantly than the others, as the values of the coefficients are not so different from each other (minimum of −0.038 for FRA:MED two-factor interaction, maximum of 0.0198 for DECONV:BS:FRA:MED four-factor interaction and mean of −0.004).

### 3.2 Enhancing and tracing the synthetic image

In order to provide more insights into the analysis of the image enhancement methods and tracing results, we used the same study procedure to test a 3D image generated synthetically. However, the synthetic image has several features that distinguish it from the image of the fungal mycelium. First, the filaments are not of uniform diameter, rather filaments near the central point from which all the filaments spread have a larger diameter, while filaments that are more distant from the origin have smaller diameters; in some cases, the diameter is reduced almost to the limit of resolution of the image (2 pixels). Second, the intensities of the pixels composing the filaments are higher than those of the fungal mycelium image, showing pixel intensities of approximately 186 compared to 106 of the image of the fungal mycelium. Finally, in the synthetic image there are no regions in which filaments are densely packed whereas most of the image of the fungal mycelium had densely packed filaments (region of the image within *y* = [0, 350]).

The synthetic image was convolved using a synthetic PSF and then noise was added, so that its quality would resemble that of a real image (see Section 2.7). In spite of the differences in the features of the two tested images, the results for SNR, SSIM, recall and precision for the synthetic image (Figure 8) were similar to those previously obtained with the image of the fungal mycelium (Figure 4). Tests 4, 6, 8 and 18 gave the highest SNR values (see the column shaded in red/pink), whereas tests 4, 18 3 and 20 had given the highest SNR values with the image of the fungal mycelium. These results confirm that the use of deconvolution and combinations of background subtraction, pixel intensity normalization and median filtering yield the greatest improvements in SNR compared to the original image (test 1). Moreover, SNR is negatively affected if Frangi vessel enhancement and median filtering are applied. As was the case for the SNR results, tests 4, 8, 6 and 18 gave the highest SSIM values. Following the same pattern as the results of the image of the fungal mycelium, the groups of tests with the highest SSIM values were 1 to 8 and 17 to 24 and the lowest values of SSIM occurred when images were enhanced with Frangi vessel enhancement followed by median filtering (tests 25-32).

**FIGURE 8.**
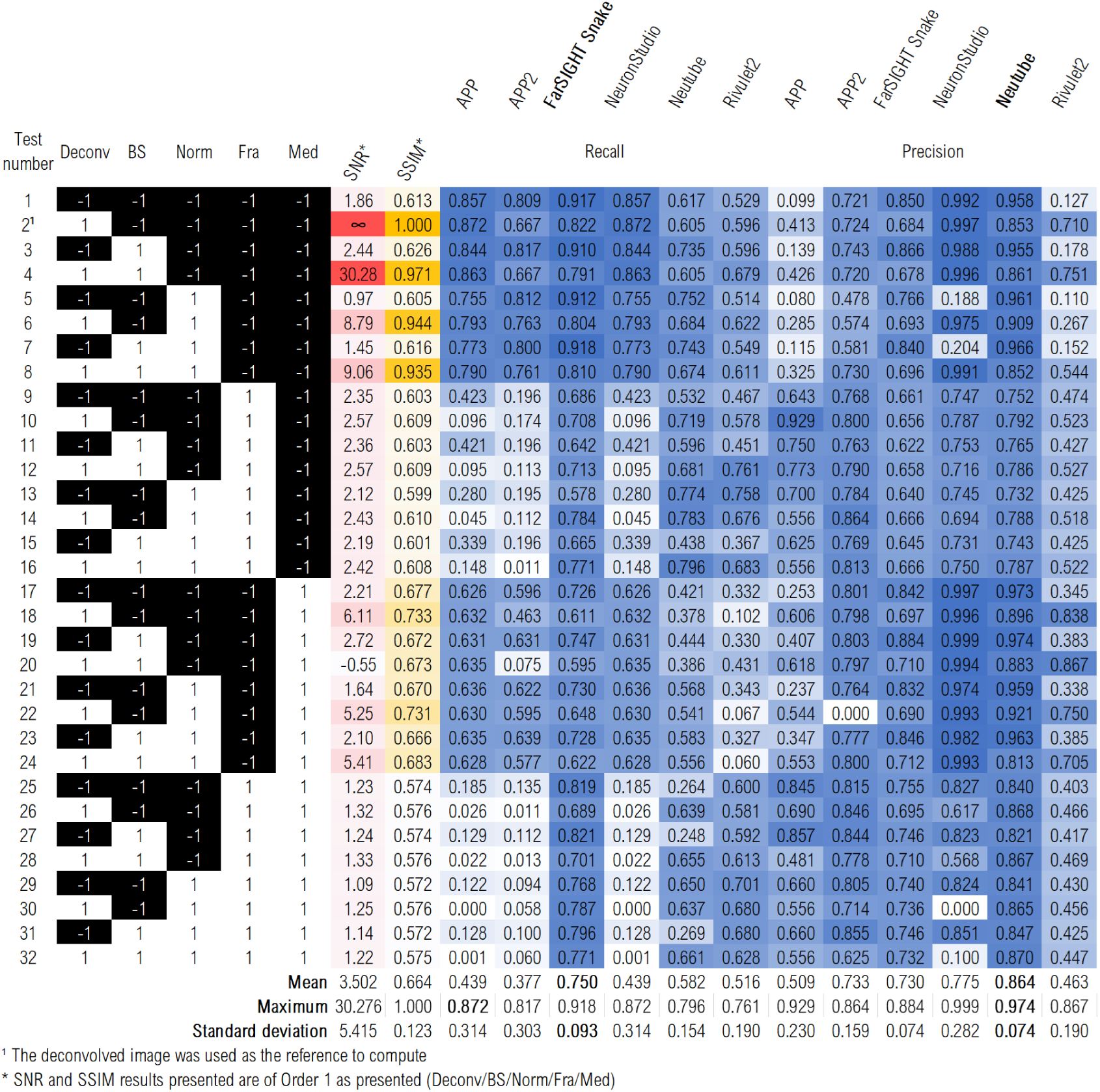
SNR, SSIM, recall and precision results of each test of the factorial design performed on the synthetic image with the order of enhancement operations *Deconv/BS/Norm/Fra/Med*.

The recall results of the synthetic image have similarities with the recall results of the image of the fungal mycelium, but with more details (i.e. no zero recall values). For instance, recall results for APP and APP2 showed drastic differences between the group of tests that used Frangi vessel enhancement (either alone or with median filtering) and the group of tests that did not apply such enhancement methods. Pixel intensity normalization affected tracing by APP in a more complex manner, as can be seen by comparing tests 1-4 with tests 5-8 and tests 17-20 with tests 21-24. It reduced recall values of APP (for instance, compare tests 1-4, which did not use pixel intensity normalization, with tests 5-8, which used it), whereas when median filtering was also applied, the negative effect of pixel intensity normalization was minimized, which indicates a synergy between these two factors. Another interesting synergic effect occurred in APP, APP2 and NeuronStudio: when deconvolution was applied without Frangi vessel enhancement, the recall results were usually higher compared to tests that did not apply deconvolution. However, when Frangi vessel enhancement was applied after deconvolution, the recall values were lower than those for the tests that did not apply deconvolution. In contrast with APP and APP2, FarSIGHT Snake and Neutube had the best overall recall values and were more robust, that is, the difference between the minimum and maximum recall values was low when the different enhancement methods were used (mean values of 0.73 and 0.582 and standard deviations of 0.093 and 0.154, respectively). However, based on the best test values, Neutube had only the 5th highest recall value, with FarSIGHT Snake (0.918), NeuronStudio and APP (both 0.872) and APP2 (0.817) having higher recall values. Furthermore, the Far-SIGHT Snake results also showed a positive synergic effect between factors: the use of Frangi vessel enhancement and median filtering increased the recall value, which did not happen in the case of the image of the fungal mycelium.

The results for the precision of the tracing methods show distinct patterns for the synthetic image. Even though the results are different in comparison with those obtained for the fungal mycelium, Rivulet2 was still the tracing method that gave the lowest precision values, with an average precision of 0.463. However, the difference among tests with high and low precision (standard deviation of 0.190) was higher than that obtained with the image of the fungal mycelium. In this manner, Rivulet2 gave both high precision (0.867 in test 20, with deconvolution, background subtraction and median filtering) and low precision (0.110 in test 5, with pixel intensity normalization). In addition, Rivulet2 did not show such a high positive influence of Frangi vessel enhancement on precision as occurred with the image of the fungal mycelium, since there was a negative synergy between Frangi vessel enhancement and median filtering. The second lowest precision values were, again, those of the APP method; the pattern of values across the various tests was similar to that obtained with the image of the fungal mycelium. The best and second best overall precision values were obtained with Neutube and NeuronStudio, with overall precision values of 0.864 and 0.775, respectively, and a standard deviation of 0.074 in both cases.

Although NeuronStudio gave the second best mean precision values (0.775), it gave the highest precision among all tracing methods with test 19 (0.999). Neutube was more sensitive to the use of Frangi vessel enhancement, with precision values that were marginally higher than those obtained in the corresponding tests with the image of the fungal mycelium. The same occurred with the use of median filtering, for which Neutube again gave marginally higher precision values than those obtained in the corresponding tests with the image of the fungal mycelium. There was also a clear positive synergic effect of deconvolution and Frangi vessel enhancement: when deconvolution was used without Frangi vessel enhancement, precision values were always lower. Tests 1-8 and 17-24 gave relatively high precision values, with test 19 giving the highest value (0.974). NeuronStudio gave clearer patterns for precision values amongst the tests than those obtained with the image of the fungal mycelium, since there were greater differences between the values (the minimum and maximum values were zero and 0.999, respectively, as opposed to 0.371 and 0.582 for the image of the fungal mycelium). Thus, despite performing better with the synthetic image, NeuronStudio was more sensitive to changes in the quality of the image. Although relatively high precision values were obtained for tests 1-8 and 17-24, these intervals also contained some relatively low values (Test 5 and 7), with these low values occurring when pixel intensity normalization was applied alone or with previous background subtraction. The lowest precision values occurred in tests 30 and 32, showing that the simultaneous use of deconvolution, pixel intensity normalization, Frangi vessel enhancement and median filtering caused NeuronStudio to fail to trace the image.

Figure 5 (b) shows the F1-score and *JSC* obtained for the various tracing methods with the synthetic image. With this image, the overall best performer was FarSIGHT Snake: it was the most robust method, with a low standard deviation of scores, and also gave the highest mean F1-score, 0.737. Neutube was the second best performer, with a mean F1-score of 0.682, but a greater spread of scores. Despite being the two best performing tracing methods overall, both FarSIGHT Snake and Neutube were outperformed by NeuronStudio when tests were evaluated individually. NeuronStudio had the best F1-scores with tests 1-4, with values up to 0.930 (test 2), whereas FarSIGHT Snake had its best F1-score, of 0.887, in test 3 and Neutube, of 0.844, in test 5. However, NeuronStudio, among all the tracing methods, gave the greatest spread of F1-scores, with a range of 0.925 (0.327 standard deviation).

Figure 7(b) shows the coefficients of the effects, both individually and in combination. Frangi vessel enhancement and median filtering had the greatest negative effects. Except for Rivulet2, the tracing methods gave lower F1-scores when Frangi vessel enhancement was applied. Also, all tracing methods gave lower F1-scores when median filtering was applied in comparison to when it was not applied. In the case of NeuronStudio, Frangi vessel enhancement alone and deconvolution followed by Frangi vessel enhancement had the greatest negative effects. Frangi vessel enhancement accounted for almost 30% of the sum of the moduli of the coefficient values, whereas deconvolution followed by Frangi vessel enhancement accounted for about 10%. However, it was advantageous to use deconvolution without Frangi vessel enhancement, as the signs of the effect coefficients changed, resulting in the highest possible F1-score in this case (test 2). The positive effects accounted for a relatively small proportion of the coefficients (24% of the sum of the moduli of the coefficients). Conversely, FarSIGHT Snake, the best overall performer and the most robust method, had smaller coefficient values and a fairly even distribution of positive and negative coefficients (positive effects accounted for about 53% of the total sum of the moduli of coefficients). Even so, only two of the main effects were positive: background subtraction and pixel intensity normalization. Additionally, the two-factor interactions of the negative effects, for example, “deconvolution and Frangi vessel enhancement” and “Frangi vessel enhancement and median filtering”, accounted for the greater part of the positive effect on the F1-score. Thus, as was the case with NeuronStudio, it was most beneficial to use a single enhancement method, background subtraction in this case, to yield the best F1-score (test 3).

## 4 DISCUSSION

The present work makes three main contributions: First, it shows that the factorial design approach is also useful to help understand the strengths and limitations of filament tracing methods, since the many enhancement operations provided a wide range of images with different qualities and features. Second, it shows that factorial designs can help researchers to evaluate the effect of image enhancement methods and choose those that fit best with their dataset. Finally, the results of this work reaffirm the importance of benchmarking filament tracing methods. In the following sections, each of the contributions will be discussed in depth.

### 4.1 Factorial designs help researchers assess the strengths and limitations of filament tracing methods

#### An assessment of strengths and limitations of the tracing methods based on their theoretical approach to tracing

Our work gives insights into the strengths of the tracing methods and allows us to assess whether the limitations that were mentioned by the authors when the tracing methods were first published are present when they are used in our test images. NeuronStudio is the oldest method available amongst the ones we tested. Our tests show that the tracing results of NeuronStudio were very poor when the enhanced images contained disconnected filaments, low intensity filaments, nonuniform intensities throughout the filaments or heavy background noise. These three features could be caused by (i) Frangi vessel enhancement, which does not enhance branch points but rather suppresses them^14;41^, (ii) any method that could reduce the foreground intensities (median filtering, Frangi vessel enhancement, deconvolution and combinations thereof) and (iii) pixel intensity normalization, which may intensify noise if applied before a noise reduction method. The fungal mycelium image used in the present work was slightly less noisy than the synthetic image (the SNR of the raw image, test 1, was 2.09 compared to the SNR of 1.86 for the synthetic image), though the fungal mycelium contained a region that had more densely packed filaments. As a result, the negative effect of pixel intensity normalization was minimized for the image of the fungal mycelium; also, the negative effect of suppressing branch points in the image was greater in the real image, which had the more complex filament tree. The high variations in the F1-score of NeuronStudio are associated with its intensity-based approach, namely voxel scooping. In voxel scooping, the image is binarized and the filament paths are traced in increments from a seed-point (in a process known as “region growing”).

APP is a graph-based tracing method. The main step in its tracing process involves the generation of a sparse graph from an oversegmented mask of the image, with vertices being foreground voxels that are connected to their direct neighbours and with edge weights that are proportional to the intensity gradient between the voxels^12^. The subsequent steps remove redundant nodes from the graph. Our tests show that APP was very sensitive to the difference between the intensities of the foreground and background: when foreground pixels were dim, APP could not detect all filaments in the image (low recall and high precision), but when foreground pixels were bright, APP resulted in overdetection of nodes (high recall and low precision). The main problem with APP comes from its oversegmentation and the existence of spurious branches in the final tracings, as observed in Figure 6(c). It appears that APP’s approach to pruning nodes that are already covered by the nodes in the centreline of the filament is not successful in situations where the diameter of the filament is greater than a few pixels. APP2 is an upgraded version of APP and has greater precision and reduced processing time^13^. It initially reconstructs the filaments with a graph-based Fast Marching algorithm, therefore reducing the size of the initial reconstruction. APP2 adds the option of generating the initial reconstruction using a grey-weighted distance transform of the image. Our results for the image of the fungal mycelium show that its use (which corresponds to the APP2-2 tests) improves the tracing results in relation to APP2 without the grey-weighted distance transform. However, the greatest improvement comes from the use of Fast Marching, as it generates a leaner initial reconstruction, which facilitates further pruning steps. Nevertheless, APP2 shares the same sensitivity to low contrast images as APP and fails to detect the entire filaments when they are dim.

FarSIGHT Snake^14;15^, which was the most robust method tested in this work, has advantages due to two main features: First, the use of Gradient Vector Flow (GVF) for both improved seed point detection and tracing (with an open active contour algorithm) and second, the implicit branch point detection. In the seed detection step, GVF is used to converge the initially detected seed points to points near the centreline of the filaments. Later, GVF is used as the snake external force of the open active contour algorithm. Despite its robustness, our results show that FarSIGHT Snake tracing was poor when filament intensities were dim. Also, FarSIGHT Snake has low precision values, due to a large number of spurious nodes in the final tracing, though this effect is smaller than in APP or Rivulet2. For instance, the number of nodes in high recall/low precision tests on the fungal mycelium image were approximately 90 · 10^3^ (test 16), 56 · 10^3^ (test 22) and 42 · 10^3^ (test 7) for Rivulet2, APP, and FarSIGHT Snake, respectively.

Neutube^19;20^ was a top performer for tests with both images. It uses a model-based approach followed by a graph-based connectivy to connect segments and resolve crossover regions. The model-based step detects filaments by fitting a 3D cylinder filter, modelled as a parameterized Laplacian of Gaussian. Then, a minimum cost spanning tree approach is used to connect segments and resolve branch points. The cost of edges between segments is calculated based on two principles: the distance between the nodes and the intensity of the voxels between the nodes. Crossovers are resolved, prior to joining segments, by computing angle changes of the end nodes of different segments (small changes in angle between two close segments will indicate that the two segments are connected). Our results show that the approach used by Neutube was robust to noise (in Figure 4 and 8, see the SNR and F1-scores in test 5: for both test images, SNR values are low but the F1-scores are high), yet sensitive to nonuniform foreground intensities (although less sensitive than NeuronStudio), dim filaments and short branches^20^. Such limitations are common to model-based (template matching of model fitting) local tracing methods such as that of Al-Kofahi *et al.*^42^.

The most recently published tracing method amongst the ones we tested is Rivulet2^43;21^. Its tracing is based on the multi-stencils fast marching method, which uses the binary distance transform of an oversegmented binary mask (with low threshold values) and further iterative back-tracing to detect branches. In our case Rivulet2 generated a huge number of nodes, therefore lowering precision values. When precision values were higher (for instance, test 18 of the synthetic image) the final tracing only covered parts of the image (the recall of 0.102 shows that only a small fraction of the filaments was traced). A reduction in recall occurred in tests where there were discontinuities in the filaments; these discontinuous filaments could not be connected by the tracing method and were later removed in the post-processing step (Rivulet2 only keeps the largest connected filament tree). However, Rivulet2 is a fast method and could be improved in case the number of nodes were reduced and the unconnected trees were kept.

A crucial point to note regarding the geometric scores we used is that the number of false positives used to calculate the F1-score and JSC is highly affected by the sampling difference between the number of nodes in the tracing result and the ground truth (Rivulet2 and APP show such a situation of a high number of false positives). Thus, it is important to visualise results carefully. For instance, upon qualitative visualisation, Rivulet2 traces the image of the fungal mycelium well, although its precision values are low due to oversegmentation. Such penalization of additional nodes could be minimized by reducing the number of nodes in the tracing graph by resampling in order to improve tracing results.

#### Connectivity analysis

The connectivity was evaluated qualitatively for the image of the fungal mycelium through visualisation of the tracing results since there was no connectivity information of a ground truth that would enable a quantitative evaluation. The visualisations showed that, for such a challenging dataset, even the best Neutube test (18) had connectivity errors, mainly crossover segments that were falsely connected. This was expected since there are regions with densely packed filaments where it is difficult to resolve whether or not they are independent segments. In addition, the filaments in the image come from more than a single source (i.e. they come from different spores) and, at this stage of fungal growth, it is impossible to determine the initial sources of the various filaments. For this reason, we also analysed the synthetic image, which is a single neuron tree with simpler connectivity. Figures 9(a-d) show that NeuronStudio (Figure 9(b)) not only had a high F1-score but also correct connectivity. As the F1-score lowered, as seen with FarSIGHT Snake (Figure 9(c)) and Neutube (Figure 9(d)), incorrect topology appeared. With FarSIGHT Snake, isolated segments that should be connected were present in the results (yellow arrows), whereas with Neutube the centre of the image, from where all filaments originate, showed incorrect connectivity (yellow arrow). Although it appears that there is a relationship between F1-scores and the connectivity, a more detailed study would be required to evaluate the connectivity of the results. For future related works, we suggest the addition of a connectivity metric such as DIADEM^22^ or the NetMets^44^ to enable a more detailed and definitive evaluation of the tracing methods based not solely on the geometrical accuracy and precision of the tracing results but also on the connectivity.

**FIGURE 9.**
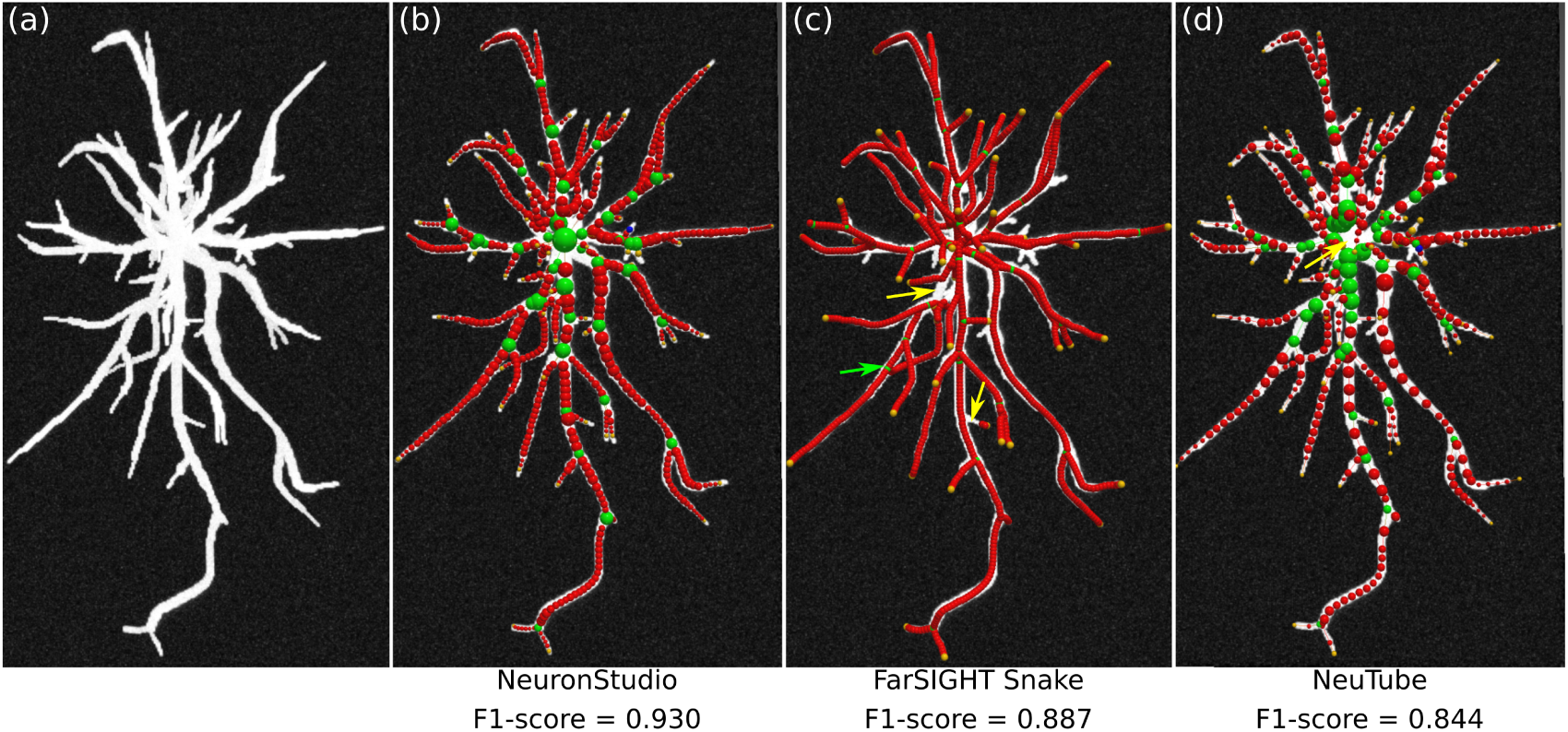
(a) Maximum intensity projection view of the synthetic image and ball-and-stick models of the best tracing results (b) NeuronStudio (test 2), (c) FarSIGHT Snake (test 3) and (d) Neutube (test 5). Red nodes correspond to body nodes, yellow nodes to end-points, green nodes to branch points and blue to seeds. The arrows show situations in which the tracing method gave incorrect topology. Yellow arrows indicate node segments that should have been connected but were not detected, whereas the green arrow shows a branch point that does not exist in the ground truth. Views were obtained with Neutube^19^

### 4.2 The factorial design approach allows for a detailed evaluation of image enhancement and filament tracing methods

#### A systematic way of visualizing effects of factors and non-additive factor interactions

Although factorial designs are commonly used to optimize outcomes of processes, our work represents the first time that a full factorial design has been used to support the evaluation of image analysis workflows. Factorial design approaches provide a more systematic way to analyse image analysis workflows, especially when several methods need to be evaluated, compared to the conventional preliminary testing and experimentation that is usually driven by the one-factor-at-a-time paradigm. Factorial design provides a mathematical model (Equation 2), the coefficients of which represent the individual effects of each factor (in this case, enhancement methods and their order in the workflow) on the image analysis outcome, as well as the effect of non-additive interactions between factors^37^. In our study, the use of a factorial design made it easier for us to identify important two-factor and three-factor interactions, for both images (Figure 7). For instance, the results of both images we tested show that the sum of the coefficients of two or more factor interactions represents more than half of the total sum of the coefficients, which indicates that these multi-factor interactions must be accounted for. In addition to indicating the existence of non-additive interaction effects, the model coefficients provide numerical values for the degree of influence of every factor and interaction on the outcome and show whether the effect is positive or negative, facilitating the interpretation of the image analysis results. For instance, a large negative coefficient value of FRA indicates that it should not be used before APP2.

Beyond that, we have proposed the use of chord graphs as an alternative to the classic Pareto plots. Chord graphs are more suitable in this situation, namely when four or more factors are investigated and a full factorial design is applied several times to test different outcomes, which, in this case, were F1-scores with different filament tracing methods. The chord graph has three advantages. First, the arc length of the chord graph allows a clear visualisation of the degree of influence of each factor on all filament tracing methods in the same graph, whereas, in the case of Pareto plots, a different plot would be necessary for each method to convey the same information. Second, the relative degree of influence of the factors for each method is more clearly shown through the widths of the chords connected to the arcs. For instance, the chord widths in the median filtering factor in Figure 7 (a) show that median filtering is an important factor affecting the tracing scores of APP and APP2, but that this factor is less important for Farsight Snake. Third, in the chord graph, the positive and negative effects are clearly distinguished through the use of opaque colour for positive effects and transparent colour for negative effects. This is the first time that a chord diagram has been used for such purpose.

#### The flexibility of factorial designs and their use for both screening important variables as well as fine optimization of the image analysis results

Our work describes the use of a full factorial design to evaluate two-level categorical factors: the use (or not) of the selected enhancement methods. However, a full factorial design could also be applied in situations where numerical parameters within the enhancement methods could be optimized or with a mixture of categorical and numerical variables. For instance, tests could be done by treating the following parameters as factors to be optimized: the standard deviation of the median filter kernel, the number of scales in which Gaussian convolution is performed in Frangi vessel enhancement or even the number of iterations and tolerance of the deconvolution process. Of course, there are other manners to perform such optimization. For example, Xu *et al.*^38^ propose the optimization of two tracing parameters through the minimization of an F-function by testing 25 different values of one parameter and 20 values of another parameter, resulting in 500 tracing tests. However, their approach may become infeasible when the range of values or the number of parameters to be optimized is higher. A factorial design approach may be more appropriate, because it uses fewer tests. For instance, if a two-factor central composite design were used in the same case presented by Xu et al.^38^, it would be possible to reduce the number of tests to multiples of 11, likely two or three times, if the starting factor levels were well selected.

Factorial designs can be used for screening or optimization^37^. In a screening design, it is common to have many factors (e.g. five or more factors in an image analysis study) to be tested and the aim is to detect the most relevant factors so that they can be further studied, but without performing too many experiments, due to time or cost constraints. Thus, screening experimental designs would be fractional factorial designs or, possibly, more specialized designs, such as the Plackett-Burman design. For example, 11 factors could be evaluated with 12 tests in a Plackett-Burman design, or a 2^6−2^ fractional factorial design could evaluate 6 two-level factors with 16 tests. In our study, we did not do a classical screening study since we evaluated the chosen factors beforehand and found them to be relevant. In addition, we identified fairly good values for the parameters within the chosen enhancement methods prior to applying the factorial design. Also, there were no time or cost constraints: each of our tests was completed in less than 10 minutes using a laptop computer (Intel® quad core processor 1.8 GHz, 16Gb RAM DDR4, GeForce MX150 4Gb graphics card) and with free and open source image analysis platforms. With 32 tests, we were able to analyse 5 factors, whereas if an approach similar to that of Xu *et al.* were used, the number of tests could easily reach hundreds of thousands of tests, thus potentially making the analysis infeasible.

In an optimization design, fewer factors are analysed, usually two or three (previously selected through screening), but with more than two levels. The additional levels are included to evaluate non-linear behaviour of the system being studied and thereby obtain a more accurate model prediction of the outcome. The central composite design is widely used for such purposes and it guides researchers towards a minimum or maximum value of the outcome within the ranges chosen for the factors considered^37;39^.

In summary, our work is a blend of both types of factorial design, for two main reasons. First, we chose to conduct a complete (full) factorial design because the chosen factors were already identified as being relevant to the tracing results and there were no time or cost constraints. Second, the tests evaluated categorical variables (the use or not of each enhancement method) with the main goal of evaluating the effect of each factor on the F1-score.

#### Usefulness of JSC, F1-score, SNR and SSIM

Researchers are usually interested in evaluating two criteria when tracing filaments: the geometric accuracy of the segmentation and the accuracy of the topology of the detected filaments (connectivity). We used the F1-score and the JSC for evaluating geometric accuracy of the detected filaments. The JSC has been previously used in a thorough comparison of particle tracking methods^23^, whereas the F1-score is commonly used to compare machine learning methods^40^, but these metrics were never used in the present context. Given spatial tolerances in the x, y and z coordinates, these values can be easily computed. The F1-score and the JSC were chosen because obtaining an accurate ground truth of the filamentous network of the fungal mycelium was made impossible by the poor quality of the image in some regions: for example, images with blurred filaments in z stacks farther away from the detector (that is, deeper within the sample, say, *z* > 50) and the existence of regions of densely packed filaments. Thus, we used the alternative approach of defining a point set ground truth in order to evaluate the accuracy of the geometric segmentation. We implemented the score computations both in Matlab® and Python and they are made available through public code repositories.

SNR is used to measure the degree of noise in an image. In our enhancement tests, we observed that some combinations of enhancement methods gave images of poorer quality (lower SNR) compared to the deconvolved image (Test 2). However, there were images with low SNR which did not show particular changes in noise levels due to the enhancement operations in relation to the deconvolved image, but had changes in the structural information in the image (for example connected filaments appeared disconnected). Thus, we also included the SSIM in our calculations, thereby evaluating changes in quality in more detail, not only with a quality parameter based on noise. In addition, we noted a counterintuitive case when either the SNR or SSIM are compared with the F1-score or other tracing scores (e.g. recall and precision): An increase in the SNR or SSIM of an enhanced image does not necessarily result in higher F1-scores, that is, we did not see any strong correlation between the SNR and the F1-score (see Supplementary Material Figures S1.5-8). This is interesting because, in image analysis, there is a common sense idea that, after an image is enhanced, it would more likely facilitate further segmentation steps and improve results. Our tests did not confirm this idea. However, we saw weak correlations between SSIM and the F1-score for some of the tracing methods, which shows that structural changes (disappearance of filament features, for example) in the image relate more easily to lower F1-score values compared to the SNR, even though such correlation is still weak. As a consequence, the use of only the SNR and SSIM to select enhancement methods is misleading. For example, for the fungal image, the highest F1-score achieved was obtained in test 18, whose SNR or SSIM values are not the highest, even though they are the third highest. Therefore, it is advisable to select enhancement methods with a complete image analysis workflow and based on geometric accuracy scores (e.g. F1 and JSC) instead of using only image quality parameters (SNR and SSIM). Ultimately, the SNR and SSIM are measurements that help in the evaluation of the result but are not as important as the F1-score.

#### The choice of the image enhancement workflow and tracing method

The chord graph in Figure 7 and the F1-scores enabled us to choose the most appropriate combination of image enhancement methods to be used prior to tracing the filaments for each of the six filament tracing methods. With the image of the fungal mycelium, the two best combinations were those of test 18 (deconvolution followed by median filtering) when Neutube was used or of test 23 (Background subtraction, followed by pixel intensity normalization and median filtering) when NeuronStudio was used. In contrast, the two best combinations for the synthetic image were those of test 2 (deconvolution) when NeuronStudio was used and of test 7 (background subtraction followed by pixel intensity normalization) when FarSIGHT Snake was used. Thus, we have not necessarily identified a general combination of enhancement methods and a tracing method that will always be optimal. The optimal combination will be image-specific, in the sense that it will depend on the quality of the original images. If the image used in the factorial design tests has a quality that is representative of the quality of the images to be processed, then one can assume that the selected combination of methods (image enhancement and tracing method) is the most appropriate. However, even when a new image to be processed does not have the same quality or filament characteristics as the tested images, our results can still guide the choice of the tracing method. This is true because we tested the tracing methods with 62 preprocessed images of quite different qualities and this allowed us to check for the robustness of the tracing methods. For instance, either FarSIGHT Snake or Neutube could be tested first in any cases, due to their robustness to changes in image quality. This robustness is attested by two results: First, FarSIGHT Snake and Neutube gave the highest mean F1-scores (0.548 and 0.685 for the image of the fungal mycelium, 0.737 and 0.682 for the synthetic image, for FarSIGHT Snake and Neutube, respectively) but also quite low standard deviations of F1-score values: the arc lengths of FarSIGHT Snake and Neutube in the chord graphs of Figure 7 show that their scores are not highly affected by the different enhancement methods used. This also leads to another conclusion: some tracing methods may require tracing tests with different image enhancement methods while other tracing methods may not require such tests. This is the case of FarSIGHT Snake and Neutube: they may give good tracing results even if the image is not enhanced.

### 4.3 Testing new tracing methods for a broad range of image qualities and filament characteristics is crucial

Although the results presented are extensive, this is not a definitive benchmark study of image enhancement methods or filament tracing methods. The analysis of the factorial design results for the F1-score show that, although similar F1-score patterns were obtained for both the image of the fungal mycelium and the synthetic image, images of different qualities and different filament features could give significant differences in tracing results. For example, the performance of APP2 was strikingly different from that of NeuronStudio. These large variations in performance would not be detected in situations where a small dataset is used. Our dataset, despite originating from only two images, was expanded because the image enhancement operations generated 62 output images of varying quality (i.e. wide range of SNR). When new tracing methods are reported, this is usually done with a smaller number of test images, although an analysis of recent papers shows that the size of test datasets is increasing. For instance, APP was tested with six images: two raw images with different filament characteristics and four where one of the raw images was processed to have different levels of noise (with the use of random bright voxel deletion)^12^. The first publication about NeuronStudio tested only one image, although it was stated that it was being used in other publications^18^. In the first Neutube paper, the tracing method was used to trace 32 neurons within a single image (filaments with the same characteristics)^19;45^. In contrast, APP2 was initially tested on the DIADEM dataset of fruitfly neurons^46^, the flycircuit.org database, the Janelia fly imagery database and other challenging datasets^13^. FarSIGHT Snake was tested on the whole DIADEM dataset^14;15^ as it was part of the DIADEM challenge and Rivulet2 was tested on both the OP dataset of DIADEM (8 images) and 114 neurons of the BigNeuron dataset^21^. Over the years, filament tracing methods have been tested more extensively, using images with different SNR and artefacts as well as images of different modalities (e.g. from confocal and brightfield microscopy)^14^. Based on our results, we encourage researchers who are developing new filament tracing methods to include not only test images of different modalities but also tests on images with different filament densities, for example, densely branched or sparse trees, and even different types of filaments, such as was done by Gonzalez et al.^47^, who included images of blood vessels, neurons and even road networks as their test datasets. Other researchers have raised similar concerns in areas where complex networks of filaments are studied, such as plant biology and neurobiology^2;14^. This way, the new methods may prove their broad applicability and stimulate discussions on their strengths and weaknesses.

## 5 CONCLUSION

In the present work, we took the challenging problem of filament tracing and evaluated different image enhancement methods and filament tracing methods through factorial designs with two images: a 3D image of a complex fungal mycelium and a 3D synthetic image of a neuronal tree. We have shown that factorial designs are powerful tools to help researchers evaluate image analysis workflows. One may choose to investigate the effects of different workflows, evaluating the results using either image quality parameters (SNR, SSIM for instance) or quantitative scores related to the image analysis problem at hand (e.g. F1-score and JSC). Regardless of the outcome chosen for the evaluation, the model that results from the factorial design gives a comprehensive analysis of the effects of the tested factors on the chosen outcome. Without an analysis of all factors simultaneously (image enhancement and tracing methods) the analysis could lead to sub-optimal results. Thus, our work gives readers an insight into the potential of the use of factorial designs in image analysis. We also identified opportunities for future extension of this work in order to explore factorial designs further and to improve the benchmarking of filament tracing methods. With respect to factorial designs, we suggest the use of a screening study followed by an optimization with other types of factorial designs, for example, the Plackett-Burman and central composite designs for screening and optimization, respectively.

Our results also show the importance of testing filament tracing workflows not only with images of different modalities, different noise and artefact levels but also with a broad range of filament characteristics (e.g. images densely populated with filaments or containing different sized filaments). If future filament tracing methods are more exhaustively tested from their conception, we believe their applicability, strengths and weaknesses may be discussed more openly and this will ensure that they are implemented and used by scientific community. Furthermore, we suggest that new benchmarking studies should include quantitative connectivity metrics to complement the analysis and provide more definitive benchmarking results.

## Supporting information

Supplemental Text and Figures (Section 1 and 2)

Supplementary spreadsheet: Parameters used for each filament tracing method

Factorial design model coefficients for the image of the fungal mycelium

Factorial design model coefficients for the synthetic image

## Acknowledgements

This research was supported by a “Universal” Grant, project number 406247/2016-1, from CNPq (Conselho Nacional de Desenvolvimento Científico e Tecnológico), a Brazilian government agency for the advancement of science and technology, and also by an ERANet-LAC Grant (ELAC2015_T03-0579), administered by CNPq through project number 443208/2016-6. Research scholarships were granted to David Mitchell and Maura Sugai-Guérios by CNPq, and to Leandro Scholz by CAPES (Coordenação de Aperfeiçoamento de Pessoal de Nível Superior), a Brazilian government agency for the development of personnel in higher education. The postgraduate program in Chemical Engineering of UFPR is financed by CAPES (Finance Code 001). The authors would like to thank members of the Laboratory of Enzyme and Fermentation Technology and the Laboratory of Enzyme Technology and Biocatalysis for the help in generating the ground-truth and Sébastien Tosi (IRB Barcelona) for helpful discussions.

## references

[1] Bonnet N. Multivariate statistical methods for the analysis of microscope image series: applications in materials science. Journal of Microscopy 1998;190(1-2):2–18. https://onlinelibrary.wiley.corn/doi/abs/10.1046/j.1365-2818.19983250876.x.

[2] Fricker MD, Moger J, Littlejohn GR, Deeks MJ. Making microscopy count: quantitative light microscopy of dynamic processes in living plants. Journal of Microscopy 2016;263(2):181–191. https://onlinelibrary.wiley.corn/doi/abs/10.1111/jrni.12403.

[3] Meijering E, Carpenter AE, Peng H, Hamprecht FA, Olivo-Marin JC. Imagining the future of bioimage analysis. Nature Biotechnology 2016 Dec;34(12):1250–1255. https://www.nature.corn/articles/nbt.3722.

[4] Scholz LA. Tracing biofilaments from images: analysis of existing methods to quantify the three-dimensional growth of filamentous fungi on solid substrates. PhD thesis, Universidade Federal do Paraná; 2018.

[5] Rubens U, Mormont R, Baecker V, Michiels G, Paavolainen L, Ball G, et al. BIAFLOWS: A collaborative framework to benchmark bioimage analysis workflows; p. 707489. https://www.biorxiv.org/content/10.1101/707489v1.

[6] Meijering E. Neuron tracing in perspective. Cytometry Part A 2010 mar;77A(7):693–704. http://doi.wiley.corn/10.1002/cyto.a.20895.

[7] Donohue DE, Ascoli GA. Automated reconstruction of neuronal morphology: An overview. Brain Research Reviews 2011;67(1):94–102.

[8] Acciai L, Soda P, Iannello G. Automated Neuron Tracing Methods: An Updated Account. Neuroinformatics 2016 Oct;14(4):353–367. http://link.springer.corn/10.1007/s12021-016-9310-0, publisher: Springer US.

[9] Wang Z, Bovik AC, Sheikh HR, Simoncelli EP. Image Quality Assessment: From Error Visibility to Structural Similarity. IEEE Transactions on Image Processing 2004 apr;13(4):600–612. http://ieeexplore.ieee.org/docurnent/1284395/.

[10] Sheppard CJR, Gan X, Gu M, Roy M. Signal-to-Noise Ratio in Confocal Microscopes. In: Handbook Of Biological Confocal Microscopy Boston, MA: Springer US; 2006.p. 442–452. http://link.springer.corn/10.1007/978-0-387-45524-2{\_}22.

[11] Sheppard CJR, Gu M, Roy M. Signal-to-noise ratio in confocal microscope systems. Journal of Microscopy 1992 ec;168(3):209–218. http://doi.wiley.corn/10.1111/j.1365-2818.1992.tb03264.x.

[12] Peng H, Long F, Myers G. Automatic 3D neuron tracing using all-path pruning. Bioinformatics 2011 jul;27(13):i239–i247.

[13] Xiao H, Peng H. APP2: automatic tracing of 3D neuron morphology based on hierarchical pruning of a gray-weighted image distance-tree. Bioinformatics 2013 jun;29(11):1448–1454.

[14] Wang Y, Narayanaswamy A, Tsai CL, Roysam B. A Broadly Applicable 3-D Neuron Tracing Method Based on Open-Curve Snake. Neuroinformatics 2011 sep;9(2-3):193–217.

[15] Narayanaswamy A, Wang Y, Roysam B. 3-D Image Pre-processing Algorithms for Improved Automated Tracing of Neuronal Arbors. Neuroinformatics 2011 sep;9(2-3):219–231.

[16] Wearne SL, Rodriguez A, Ehlenberger DB, Rocher AB, Henderson SC, Hof PR. New techniques for imaging, digitization and analysis of three-dimensional neural morphology on multiple scales. Neuroscience 2005 jan;136(3):661–680.

[17] Rodriguez A, Ehlenberger DB, Dickstein DL, Hof PR, Wearne SL. Automated Three-Dimensional Detection and Shape Classification of Dendritic Spines from Fluorescence Microscopy Images. PLoS ONE 2008 apr;3(4):1–12.

[18] Rodriguez A, Ehlenberger DB, Hof PR, Wearne SL. Three-dimensional neuron tracing by voxel scooping. Journal of Neuroscience Methods 2009 oct;184(1):169–175.

[19] Feng L, Zhao T, Kim J. neuTube 1.0: a New Design for Efficient Neuron Reconstruction Software Based on the SWC Format. eNeuro 2015;http://www.eneuro.org/content/early/2015/01/02/ENEUR0.0049-14.2014.

[20] Zhao T, Xie J, Amat F, Clack N, Ahammad P, Peng H, et al. Automated Reconstruction of Neuronal Morphology Based on Local Geometrical and Global Structural Models. Neuroinformatics 2011 sep;9(2-3):247–261.

[21] Liu S, Zhang D, Song Y, Peng H, Cai W. Automated 3D Neuron Tracing with Precise Branch Erasing and Confidence Controlled Back-Tracking. IEEE Transactions on Medical Imaging 2018;p. 1–1. https://ieeexplore.ieee.org/docurnent/8354803/.

[22] Gillette TA, Brown KM, Ascoli GA. The DIADEM metric: comparing multiple reconstructions of the same neuron. Neuroinformatics 2011 sep;9(2-3):233–45. http://www.ncbi.nlrn.nih.gov/pubrned/21519813http://www.pubrnedcentralnih.gov/articlerender.fcgi?artid=PMC4339018.

[23] Chenouard N, Smal I, de Chaumont F, Maška M, Sbalzarini IF, Gong Y, et al. Objective comparison of particle tracking methods. Nature Methods 2014 jan;11(3):281–289. http://www.nature.corn/doifinder/10.1038/nrneth.2808.

[24] Sugai-Guérios MH. Understanding the growth of hyphae of filamentous fungi on the surfaces of solid media through computational models and confocal microscopy. PhD thesis, Federal University of Santa Catarina; 2016.

[25] Kirshner H, Aguer F, Sage D, Unser M. 3-D PSF fitting for fluorescence microscopy: implementation and localization application. Journal of Microscopy 2013 jan;249(1):13–25. http://doi.wiley.corn/10.1111/j.1365-2818.2012.03675.x.

[26] Sage D, Donati L, Soulez F, Fortun D, Schmit G, Seitz A, et al. DeconvolutionLab2: An open-source software for deconvolution microscopy. Methods 2017 feb;115:28–41. https://www.sciencedirect.corn/science/article/pii/S1046202316305096?via%3Dihub.

[27] Sternberg SR. Biomedical Image Processing. Computer 1983 jan;16(1):22–34. http://ieeexplore.ieee.org/docurnent/1654163/.

[28] Frangi AF, Niessen WJ, Vincken KL, Viergever MA. Multiscale vessel enhancement filtering. In: Wells WM, Colch-Ester A, Delp S, editors. Medical Image Computing and Computer-Assisted Intervention — MICCAI’98. MICCAI 1998. Lecture Notes in Computer Science, vol 1496 Springer Berlin Heidelberg; 1998.p. 130–137. http://link.springer.corn/10.1007/BFb0056195.

[29] Schindelin J, Arganda-Carreras I, Frise E, Kaynig V, Longair M, Pietzsch T, et al. Fiji: an open-source platform for biological-image analysis. Nature Methods 2012 jul;9(7):676–682. http://www.nature.corn/articles/nrneth.2019.

[30] Castle M, Keller J, Rolling Ball Background Subtraction implementation; 2007. http://irnagej.net/plugins/rolling-ball.htrnl.

[31] Cuntz H, Forstner F, Borst A, Häusser M. One Rule to Grow Them All: A General Theory of Neuronal Branching and Its Practical Application. PLOS Computational Biology 2010 08;6(8):1–14. https://doi.org/10.1371/journal.pcbi1000877.

[32] Peng H, Ruan Z, Atasoy D, Sternson S. Automatic reconstruction of 3D neuron structures using a graph-augmented deformable model. Bioinformatics 2010 jun;26(12):i38–i46.

[33] Schindelin J, Meijering E, RandomJ; 2014. https://irnagescience.org/rneijering/software/randornj/

[34] Dinno A, Package ‘dunn.test’; 2017. https://cran.rstudio.corn/web/packages/dunn.test/dunn.test.pdf

[35] Peng H, Ruan Z, Long F, Simpson JH, Myers EW. V3D enables real-time 3D visualization and quantitative analysis of large-scale biological image data sets. Nature Biotechnology 2010 apr;28(4):348–353. http://www.nature.corn/doifinder/10.1038/nbt.1612.

[36] Gu Z, Gu L, Eils R, Schlesner M, Brors B. circlize implements and enhances circular visualization in R. Bioinformatics 2014 Oct;30(19):2811–2812. https://acadernic.oup.corn/bioinforrnatics/article-lookup/doi/10.1093/bioinforrnatics/btu393.

[37] Box GEP, Hunter JS, Hunter WG. Statistics for experimenters: design, innovation, and discovery. 2 ed. Wiley series in probability and statistics, Wiley-Interscience; 2005.

[38] Xu T, Vavylonis D, Tsai FC, Koenderink GH, Nie W, Yusuf E, et al. SOAX: A software for quantification of 3D biopolymer networks. Scientific Reports 2015 mar;5:9081.

[39] Myers RH, Montgomery DC. Response surface methodology: process and product optimization using designed experiments. Wiley series in probability and statistics: Applied probability and statistics, Wiley; 1995. https://books.googlecorn.br/books?id=7xvvAAAAMAAJ.

[40] Caicedo JC, Roth J, Goodman A, Becker T, Karhohs KW, Broisin M, et al. Evaluation of Deep Learning Strategies for Nucleus Segmentation in Fluorescence Images. bioRxiv 2019;https://www.biorxiv.org/content/early/2019/02/06/335216.

[41] Obara B, Fricker M, Gavaghan D, Grau V. Contrast-Independent Curvilinear Structure Detection in Biomedical Images. IEEE Transactions on Image Processing 2012 May;21(5):2572–2581. http://ieeexplore.ieee.org/docurnent/6140570/.

[42] Al-Kofahi KA, Lasek S, Szarowski DH, Pace CJ, Nagy G, Turner JN, et al. Rapid automated three-dimensional tracing of neurons from confocal image stacks. IEEE Transactions on Information Technology in Biomedicine 2002 jun;6(2):171–187.

[43] Liu S, Zhang D, Liu S, Feng D, Peng H, Cai W. Rivulet: 3D Neuron Morphology Tracing with Iterative Back-Tracking. Neuroinformatics 2016 oct;14(4):387–401. http://link.springer.corn/10.1007/s12021-016-9302-0.

[44] Mayerich D, Bjornsson C, Taylor J, Roysam B. NetMets: software for quantifying and visualizing errors in biological network segmentation. BMC Bioinformatics 2012 13:8 2012;13(8):1–19.

[45] Druckmann S, Feng L, Lee B, Yook C, Zhao T, Magee JC, et al. Structured Synaptic Connectivity between Hippocampal Regions. Neuron 2014 Feb;81(3):629–640. http://www.sciencedirect.corn/science/article/pii/S0896627313010945.

[46] Brown KM, Barrionuevo G, Canty AJ, De Paola V, Hirsch JA, Jefferis GS, et al. The DIADEM data sets: representative light microscopy images of neuronal morphology to advance automation of digital reconstructions. Neuroinformatics 2011;9(2-3):143–157.

[47] Gonzalez G, Fleurety F, Fua P. Learning rotational features for filament detection. In: 2009 IEEE Conference on Computer Vision and Pattern Recognition IEEE; 2009. p. 1582–1589.

