## Supplemental Text and Figures (Section 1 and 2) for "Factorial design as a tool to evaluate image analysis workflows systematically: its application to the filament tracing problem"

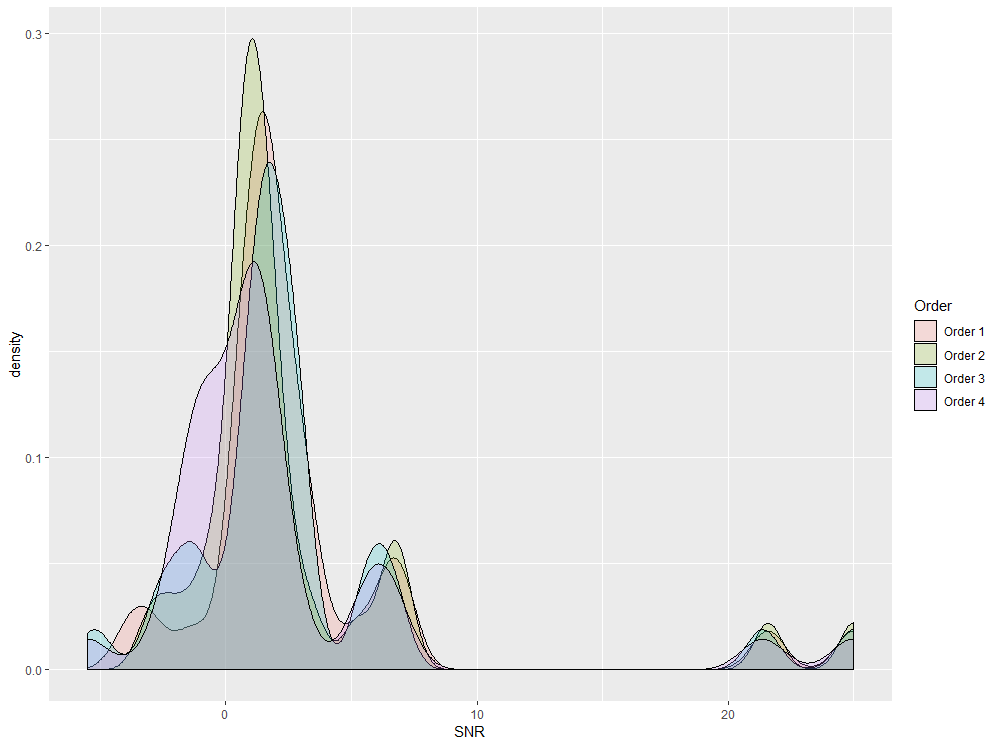

Figure S1.1 - Density plot of the SNR of the test image of the fungal mycelium for the four different orders of enhancement operations, as per Table 1.

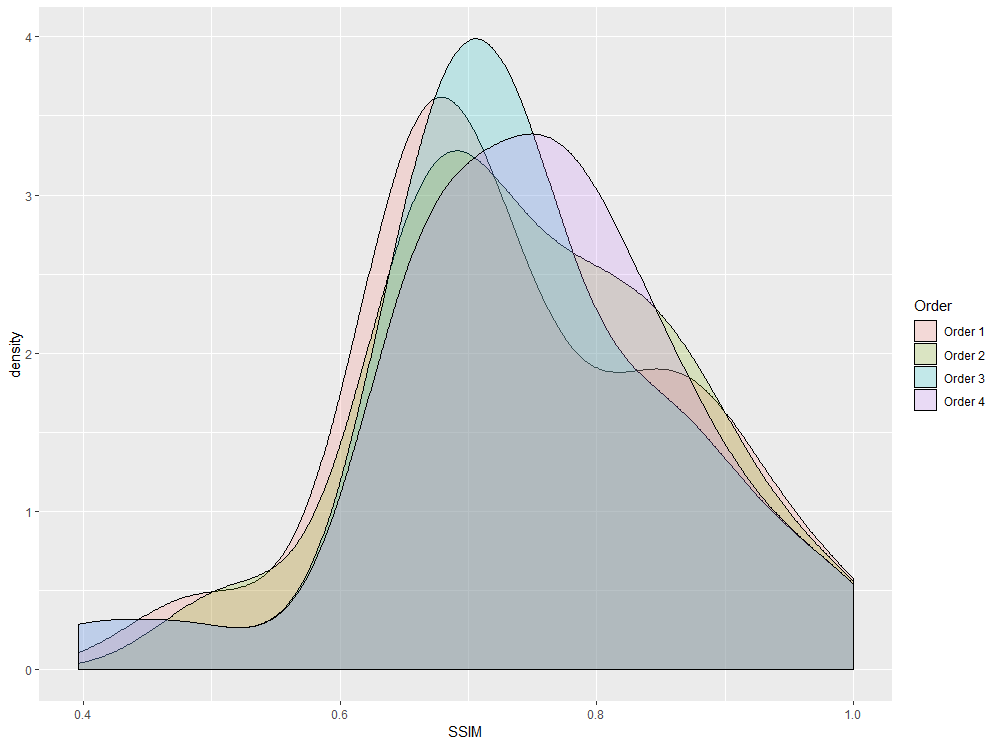

Figure S1.2 - Density plot of the SSIM of the test image of the fungal mycelium for the four different orders of enhancement operations, as per Table 1.

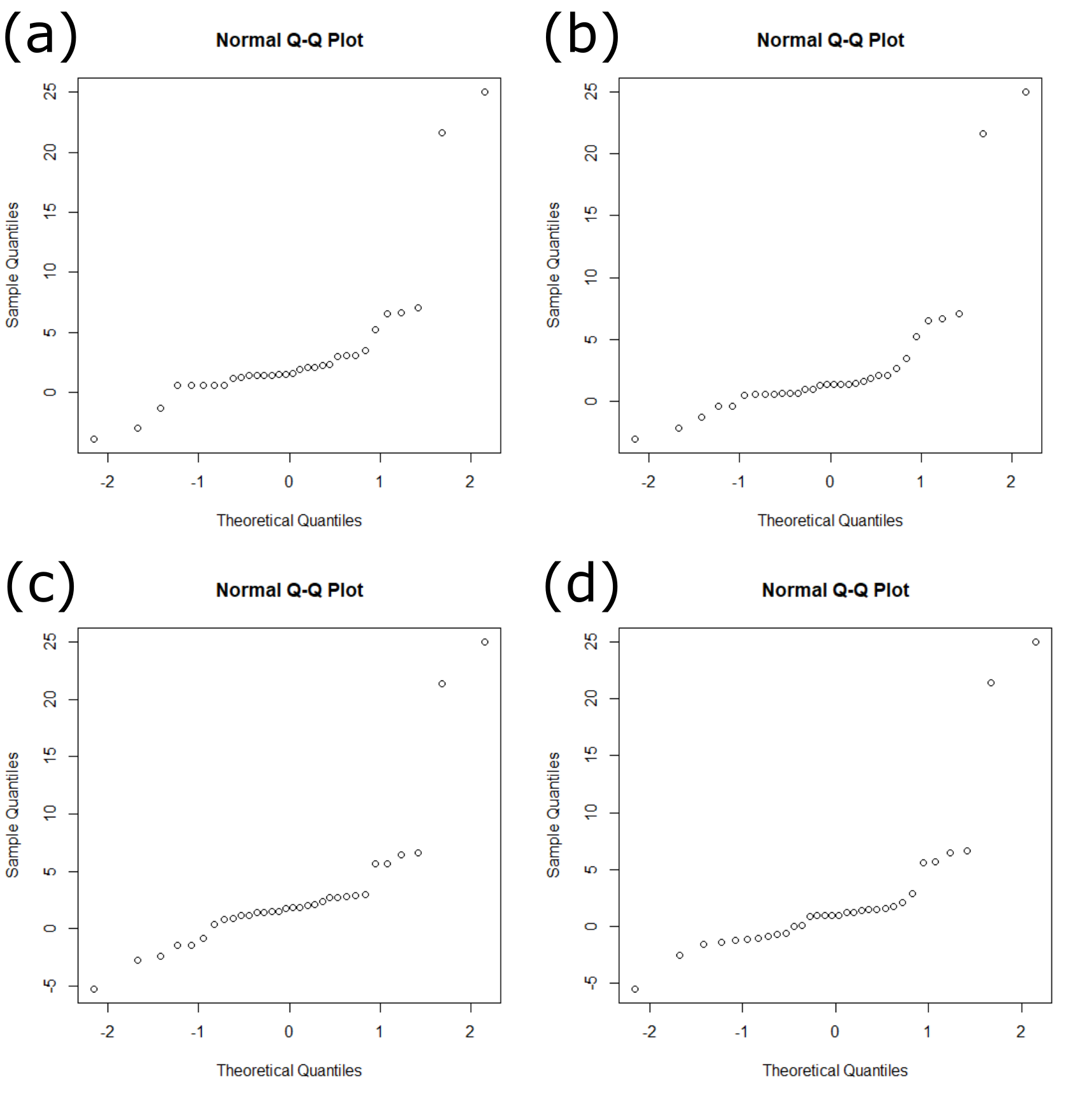

Figure S1.3 – Normal Q-Q plots of SNR of the test images of the fungal mycelium. From top left to bottom right, Orders (a) 1, (b) 2, (c) 3 and (d) 4 are shown.

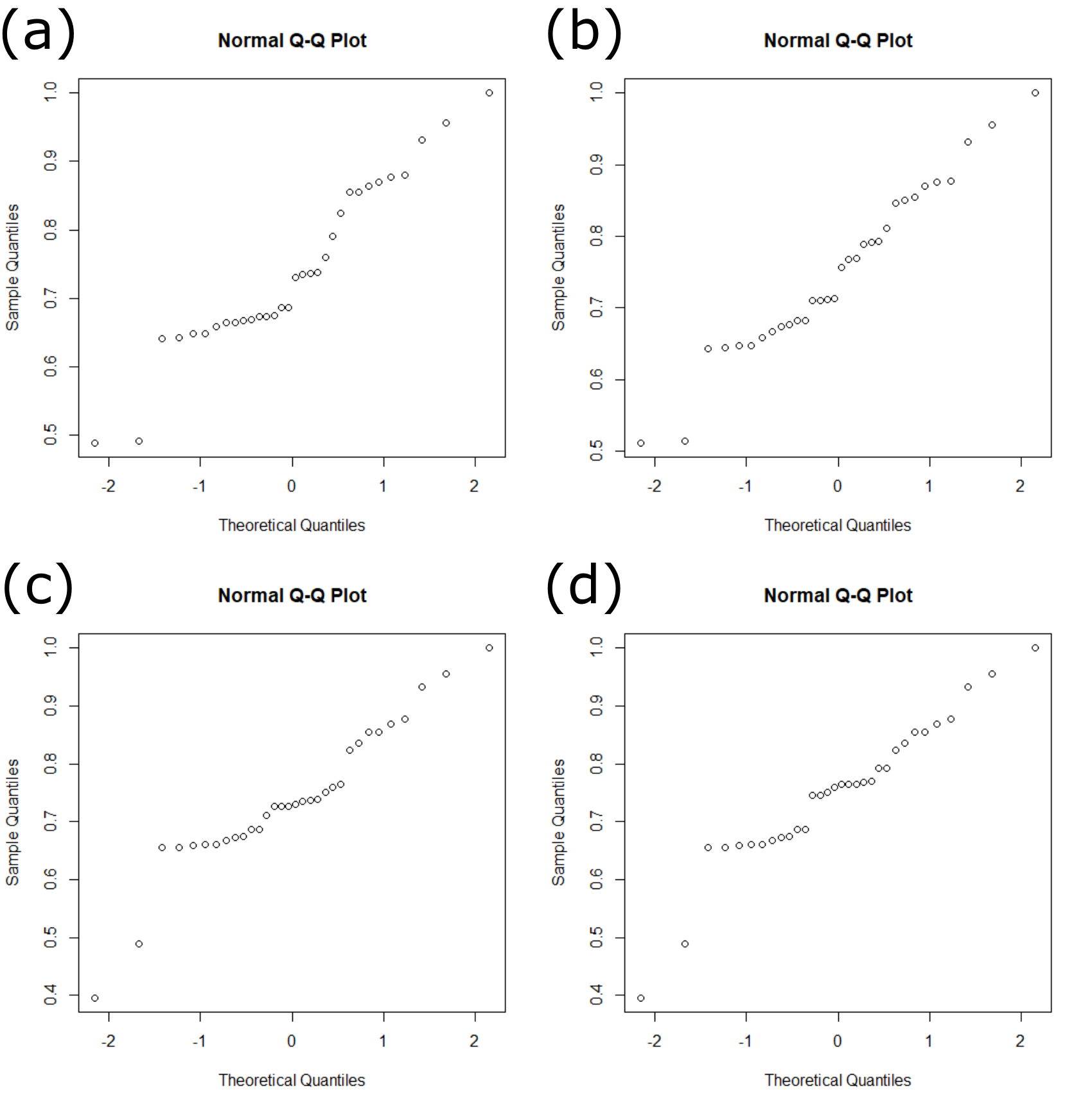

Figure S1.4 – Normal Q-Q plots of SSIM of the test images of the fungal mycelium. From top left to bottom right, Orders (a) 1, (b) 2, (c) 3 and (d) 4 are shown.

Table S1.1 – Statistical tests to verify normality of the dataset and Analysis of Variance (ANOVA) and equivalent non-parametric Kruskal-Wallis tests.

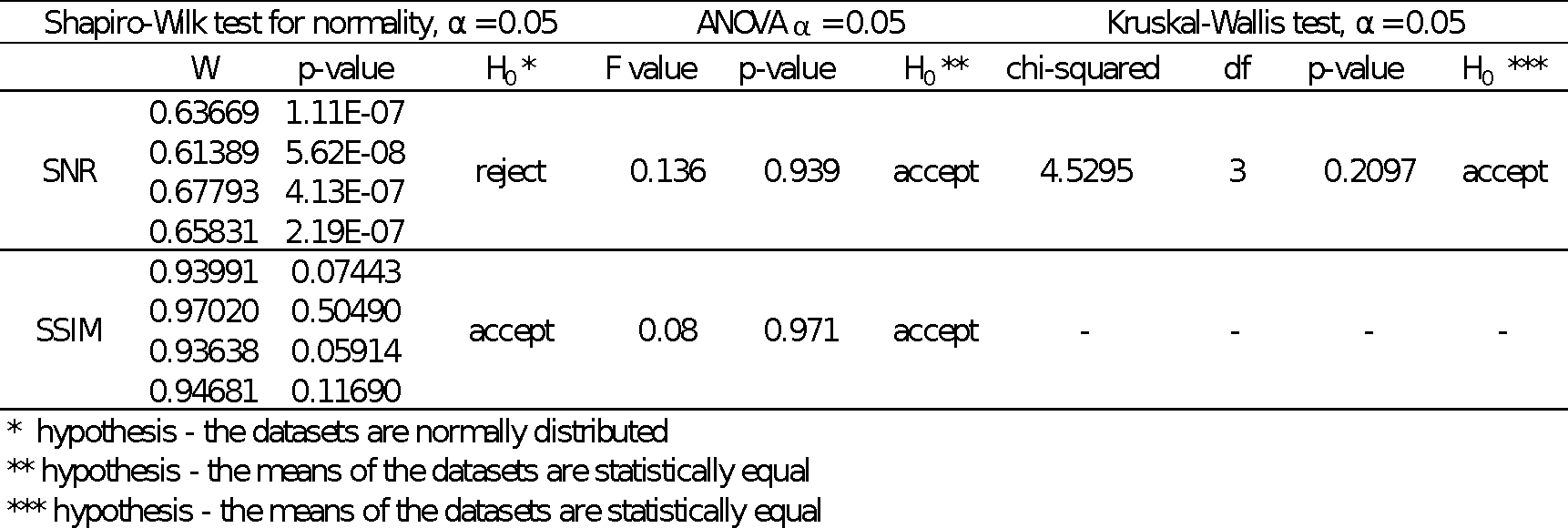

Table S1.2 F1-score results of the tests of the factorial design for the image of the fungal mycelium

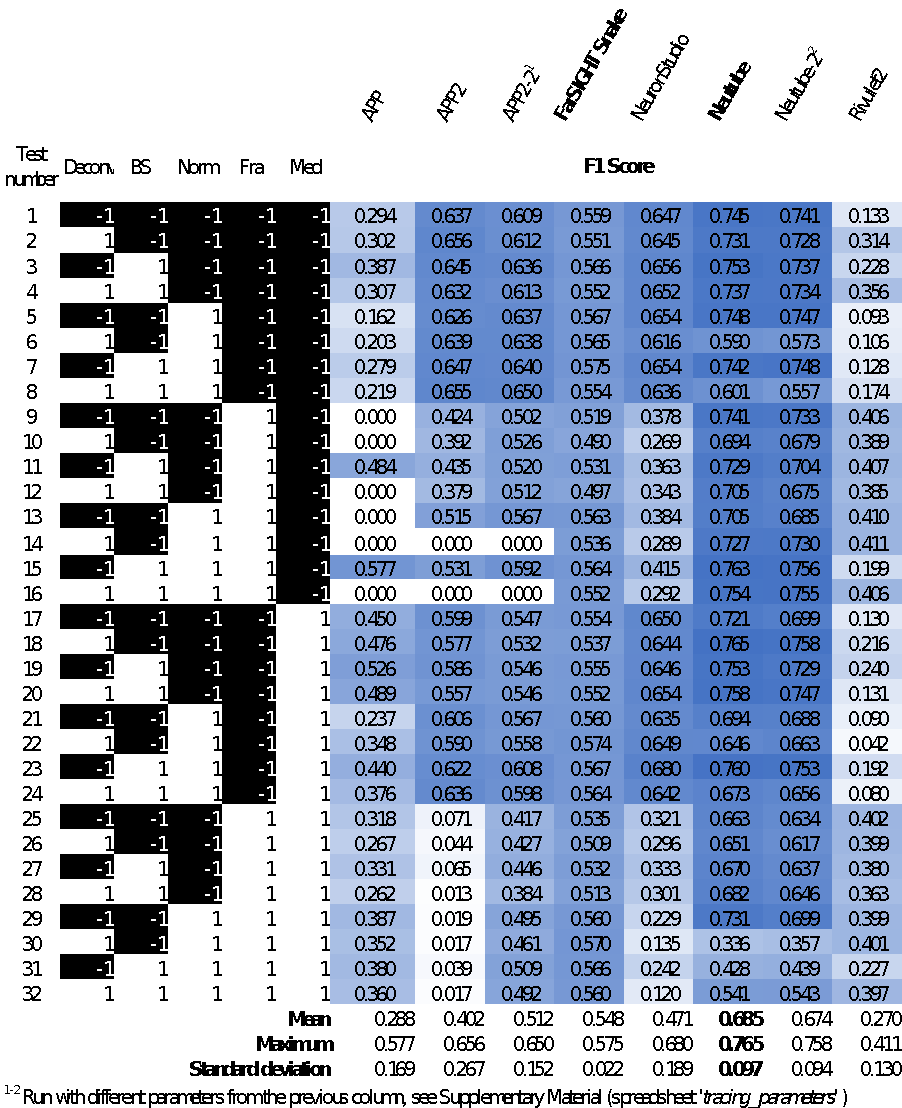

Table S1.2 F1-score results of the tests of the factorial design for the synthetic image of a neuronal tree

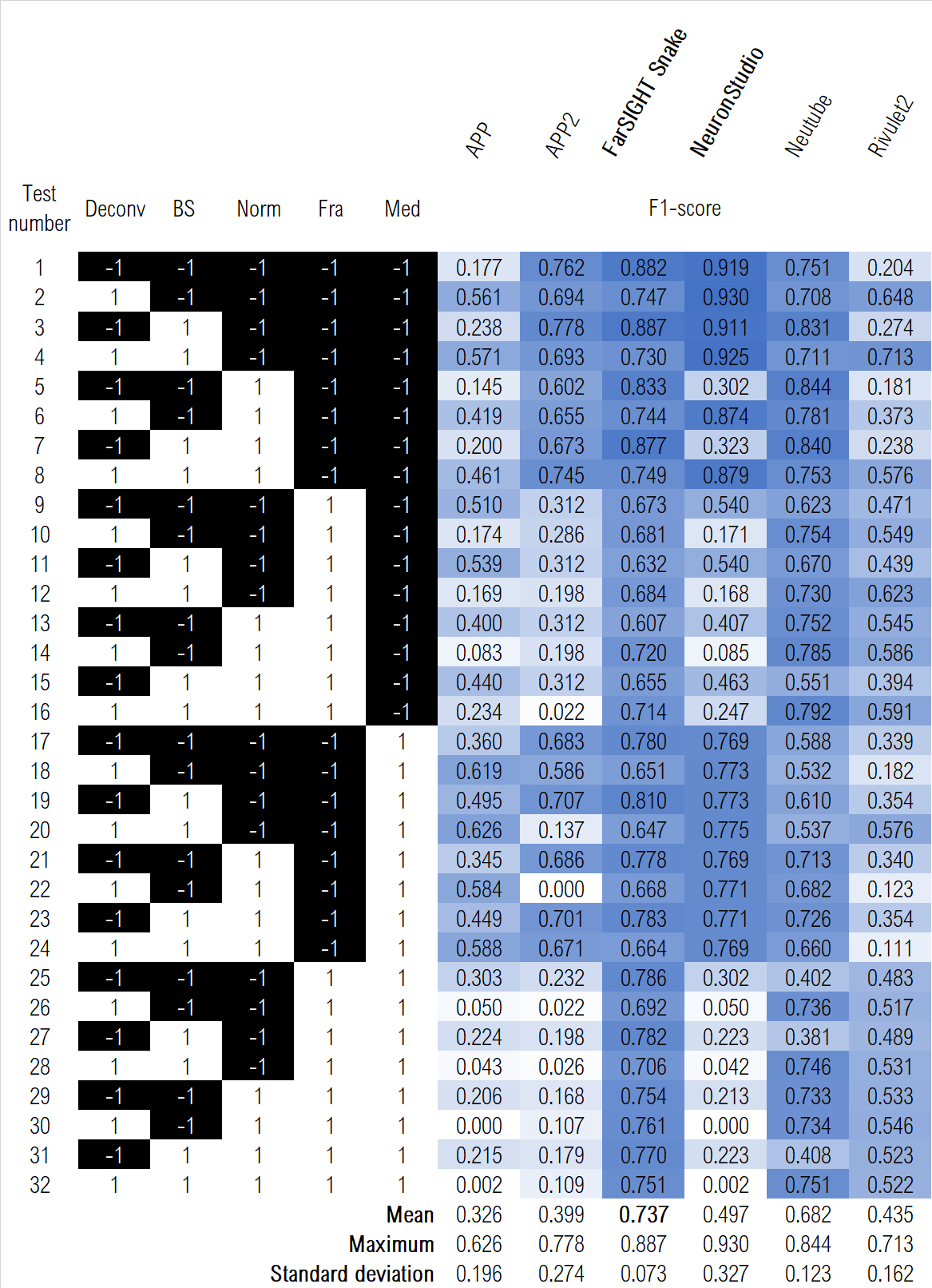

**Relationship between SNR or SSIM and F1-scores**

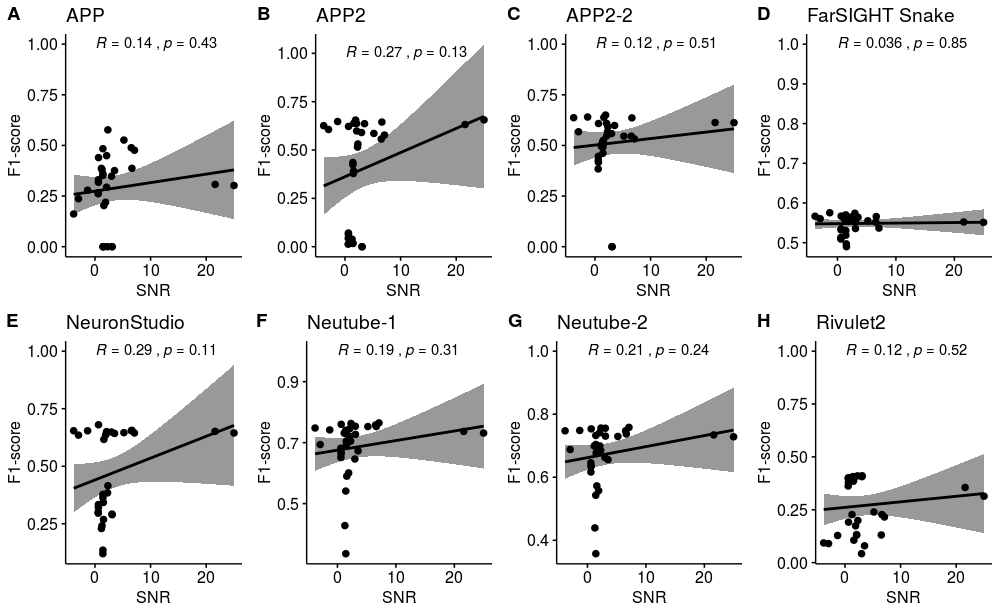

Figure S1.5 (A-H) Correlation plots, with corresponding Pearson correlation coefficients (R) and p-values (α = 0.05), of the relationship between SNR and F1-score for the image of the fungal mycelium.

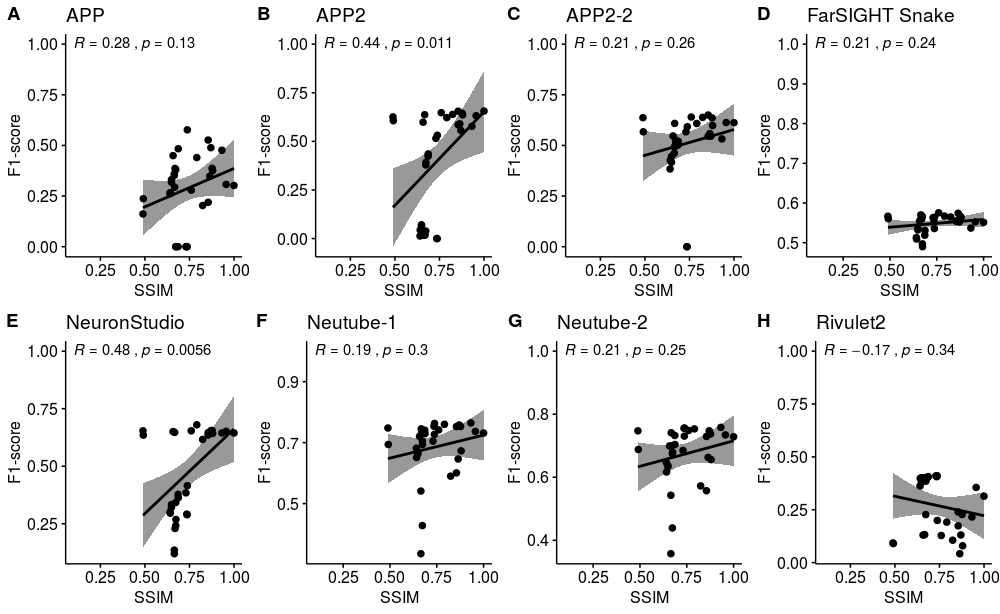

Figure S1.6 (A-H) Correlation plots, with corresponding Pearson correlation coefficients (R) and p-values (α = 0.05), of the relationship between SSIM and F1-score for the image of the fungal mycelium.

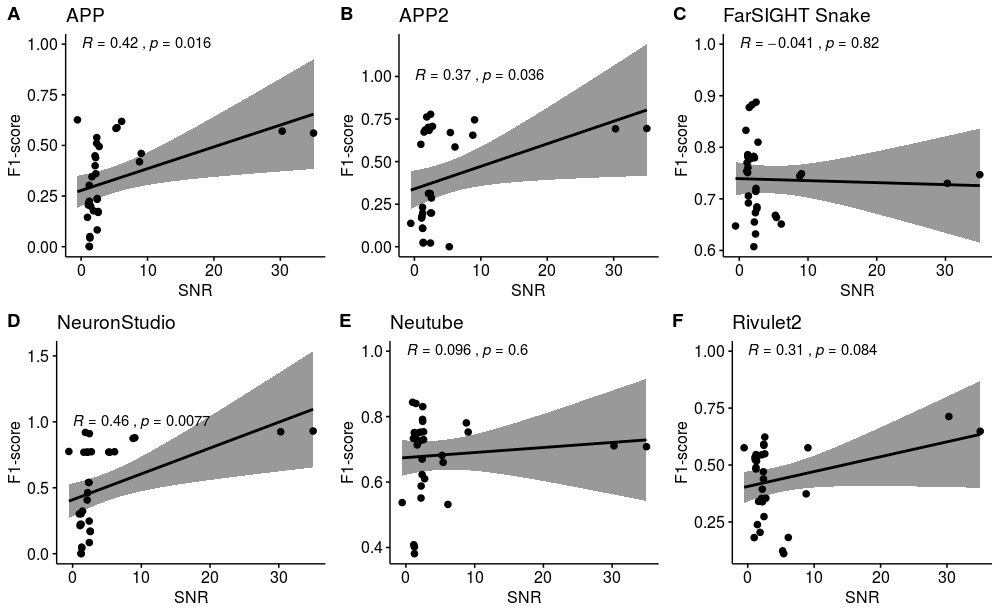

Figure S1.7 (A-F) Correlation plots, with corresponding Pearson correlation coefficients (R) and p-values (α = 0.05), of the relationship between SNR and F1-score for the synthetic image of a neuronal tree.

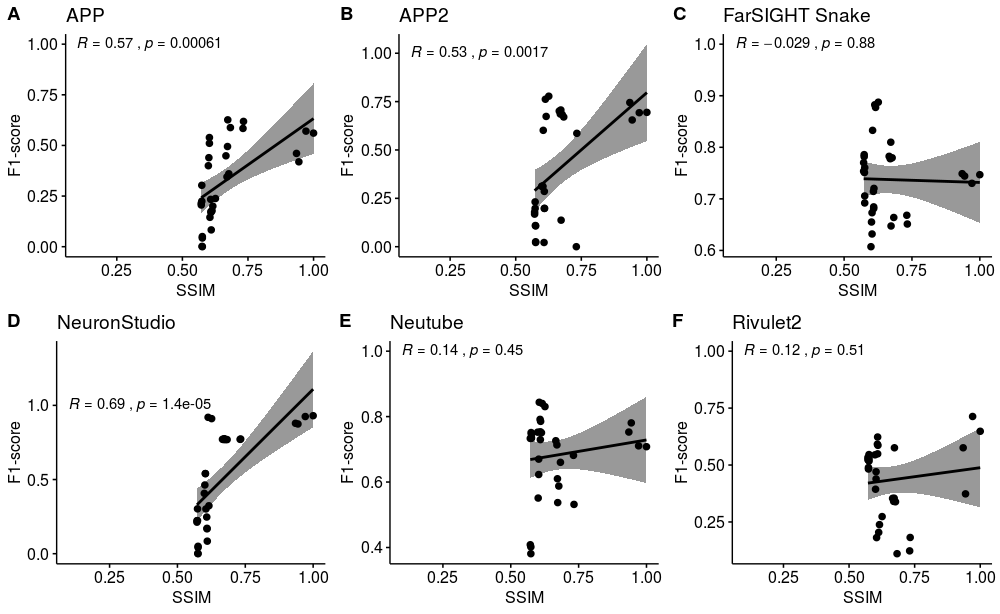

Figure S1.8 (A-F) Correlation plots, with corresponding Pearson correlation coefficients (R) and p-values (α = 0.05), of the relationship between SSIM and F1-score for the synthetic image of a neuronal tree.

In our work did not compute tracing scores for the other orders of enhancement operations (2, 3 and 4). However, since we found that there was no clear correlation between SNR and the F1-score and only a weak correlation between SSIM and F1-score, it could be possible that the order of enhancement operations could give better F1-score results, even if we observed that there is no statistical difference between the SNR and SSIM score result among the orders of enhancement operations. This is to be further studied in future works.

### Section 2

**Procedure to obtain ground truth annotations to evaluate filament tracing methods**

This procedure describes the steps to annotate data points, which will be used as ground truth, from the image of the fungal mycelium. These data points will be used to evaluate the results of filament tracing methods.

1. **Open ImageJ**

Open Fiji ImageJ on your operating system of preference. Fiji window should look similar to this:

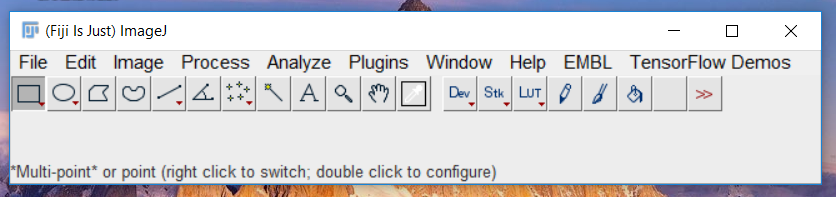

2. **Open the image you wish to annotate**

There are two ways to open an image. The first option is to go to *File > Open* and select the path and the file you wish to open. The second is to simply drag and drop the file to Fiji window from an open file explorer.

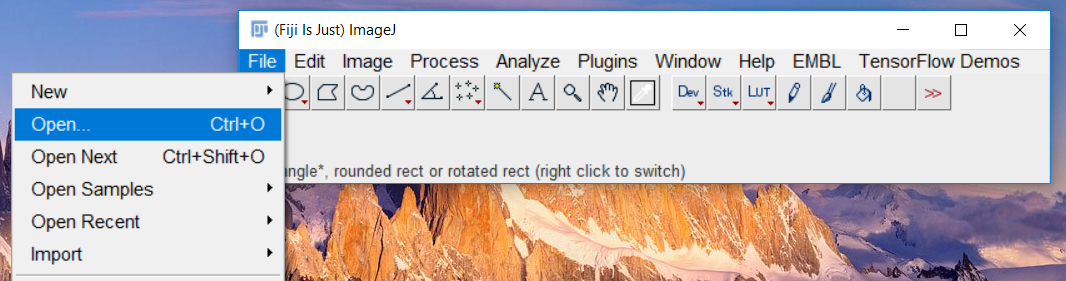

In our case, the images will be similar to the one shown below:

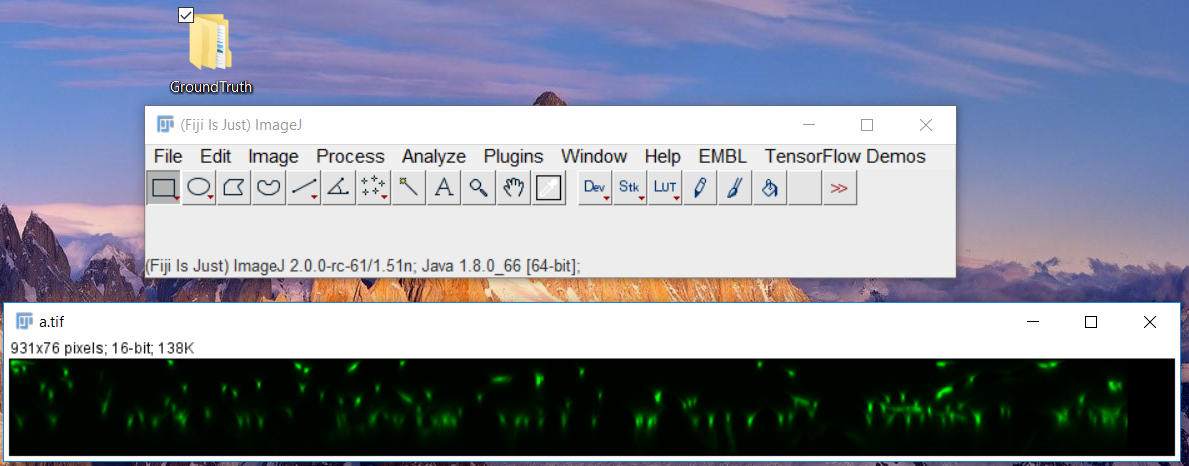

3. **Use multi-point tool to annotate the image**

In order to have a set of points that will be used to construct the ground truth, we use the multi-point tool. In order to activate the multi-point tool, you should click on one of the following buttons.

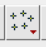
*Multi-point Tool*
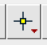
*Point Tool*

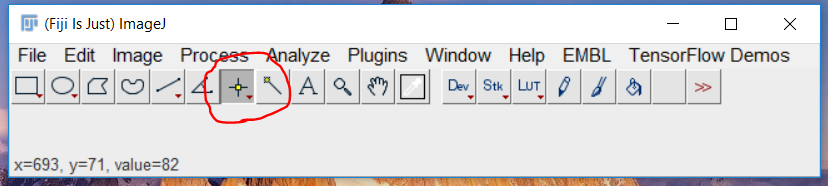

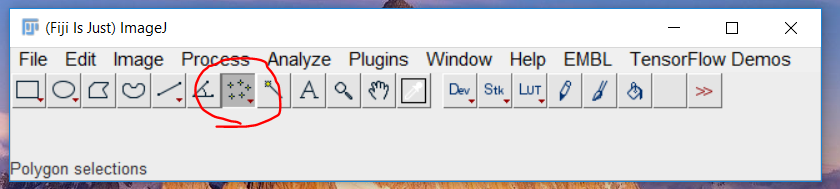

In case Fiji shows *Point Tool* instead of the *Multi-Point tool*, right click on the button and select ***Multi-Point tool*** as shown below.

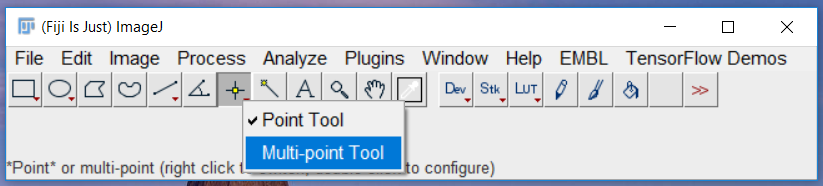

4. **Annotate the image based on defined criteria.**
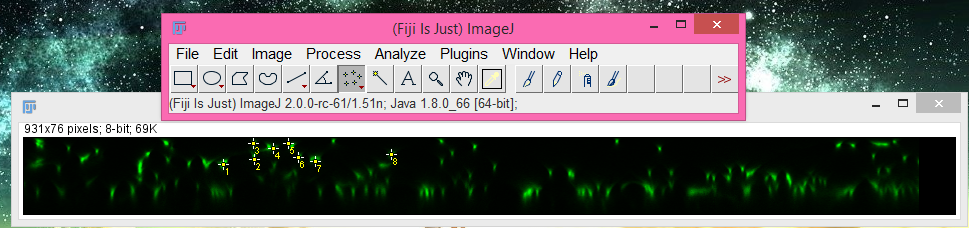

In this case, the filament segments we wish to annotate will most often appear as blobs and not curvilinear (tubular) structures. Thus, select the center of the blob where there appears to be a peak in intensity.

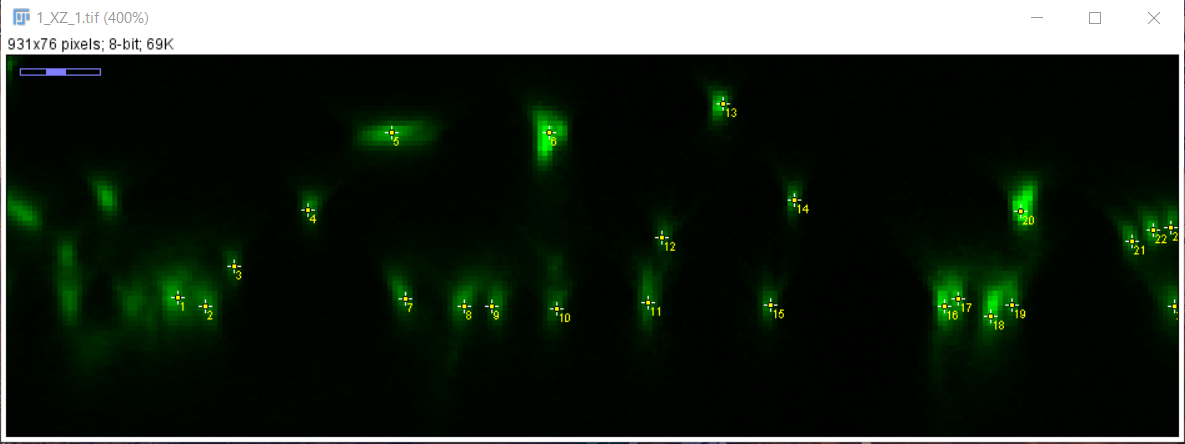

In order to zoom in and out of the image and facilitate the visualisation of the regions being annotated, use *ctrl + ‘+’* or *ctrl + ‘-’* (Image> Zoom > In/Out) or even *ctrl + mouse scroller up/down*. To navigate through the image when it is zoomed, use the *scroll tool* (the ‘hand’). In case you are using a mouse you can use the mouse scroll button (at least when using Windows and Linux Ubuntu).

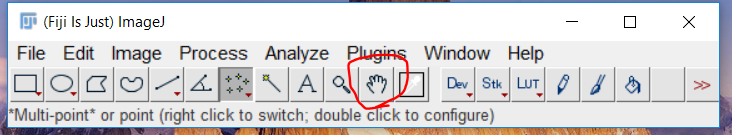

5. **Extract the annotated points as a table**

After all the points the annotator deems to be correct are selected, the set of data points are extracted. With the image selected, go to *Analyze > Measure*.

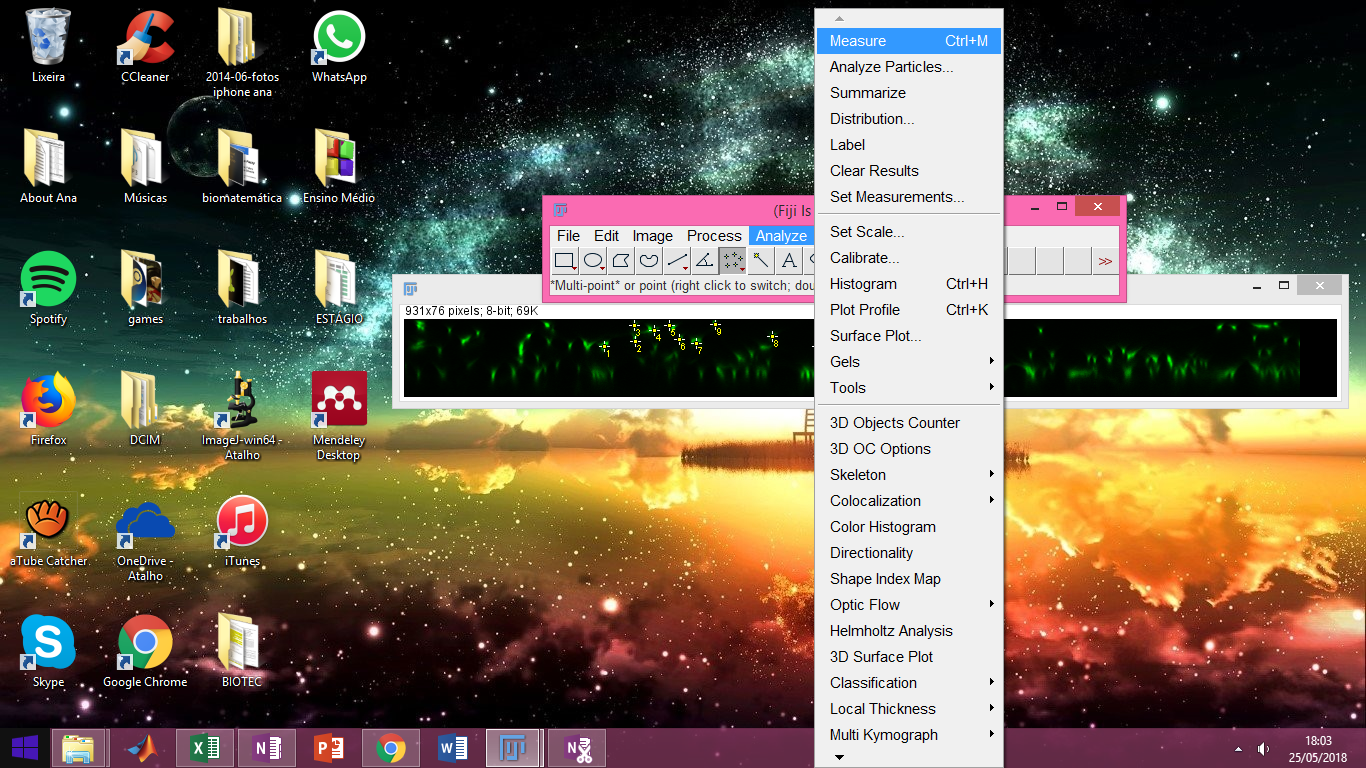

After selecting *Measure*, a new window will appear with the results shown in a table.

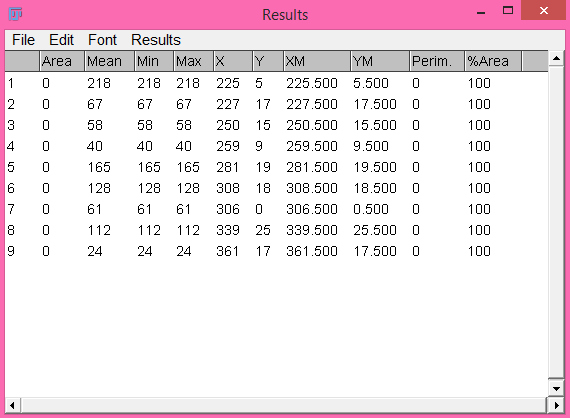

This is the list that gives information about the all the annotated points. The only mandatory information is the coordinate values of the position of each point (X and Y columns).

**6. Save results**

In the Results window, go to *File > Save*. Save the document with a naming pattern of your choice. Later this naming pattern will be used in the Matlab code to determine the ground truth points (Mean cluster points). For this work, we used the following naming pattern: “Results_*imagename*_*yourname*”. For instance, if Leandro annotated image “2_XZ_21” , the results file name should be “Results_2_XZ_21_Leandro”.

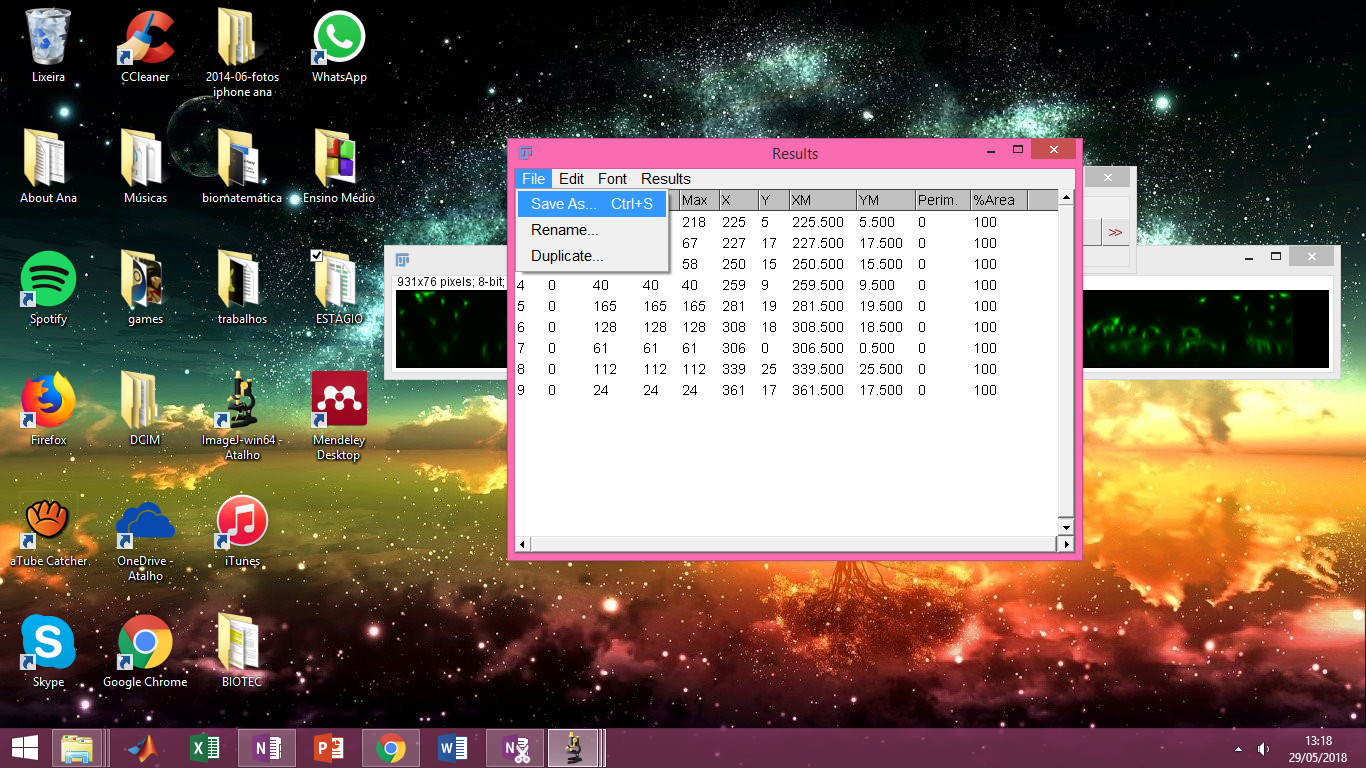
